# Discovery and functional assessment of a novel adipocyte population driven by intracellular Wnt/β-catenin signaling in mammals

**DOI:** 10.1101/2021.01.14.426702

**Authors:** Zhi Liu, Tian Chen, Sicheng Zhang, Tianfang Yang, Yun Gong, Hong-Wen Deng, Ding Bai, Weidong Tian, YiPing Chen

**Affiliations:** Department of Cell and Molecular Biology, Tulane University, New Orleans, LA, USA; State Key Laboratory of Oral Diseases, National Clinical Research Center for Oral Disease, West China Hospital of Stomatology, Sichuan University, Chengdu, Sichuan Province, China; Tulane Center of Biomedical Informatics and Genomic, Deming Department of Medicine, School of Medicine, Tulane University, New Orleans, LA, USA

## Abstract

Wnt/β-catenin signaling has been well established as a potent inhibitor of adipogenesis. Here, we identified a population of adipocytes that exhibit persistent activity of Wnt/β-catenin signaling, as revealed by the *Tcf*/*Lef-Gfp* reporter allele, in embryonic and adult mouse fat depots, named as Wnt^+^ adipocytes. We showed that the β-catenin-mediated signaling activation in these cells is Wnt ligand- and receptor-independent but relies on AKT/mTOR pathway and is essential for cell survival. Such adipocytes are distinct from classical ones in transcriptomic and genomic signatures and can be induced from various sources of mesenchymal stromal cells including human cells. Genetic lineage-tracing and targeted cell ablation studies revealed that these adipocytes convert into beige adipocytes directly and are also required for beige fat recruitment under thermal challenge, demonstrating both cell autonomous and non-cell autonomous roles in adaptive thermogenesis. Furthermore, mice bearing targeted ablation of these adipocytes exhibited glucose intolerance, while mice receiving exogenously supplied such cells manifested enhanced glucose utilization. Our studies uncover a unique adipocyte population in regulating beiging in adipose tissues and systemic glucose homeostasis.

## INTRODUCTION

Adipose tissues, consisting of white adipose tissue (WAT) and brown adipose tissue (BAT), play a critical role in maintaining whole-body metabolic homeostasis, with WAT serving for energy storage and BAT for energy dissipation to produce heat (Cannon and Nedergaard, 2004; Rosen and Spiegelman, 2014). In addition to white and brown fat cells that are differentiated from heterogenous stromal vascular fractions (SVFs) of distinct lineages (Hepler et al., 2017; Rosenwald and Wolfrum, 2014; Schwalie et al., 2018), a third type of inducible adipocyte exists known as beige or brite adipocytes that are transiently generated in WAT depots in response to external stimulations such as environmental cold acclimation (Bostrom et al., 2012; Ikeda et al., 2018; Wu et al., 2012). Like brown adipocytes, activated beige adipocytes through ‘‘beiging’’ or ‘‘browning’’ process of WAT express key thermogenic marker uncoupling protein 1 (UCP1) and exert adaptive thermogenic function (Ishibashi and Seale, 2010; Petrovic et al., 2010). Despite that the capacity for thermogenic fat cells (i.e., brown and beige adipocytes) to protect against diet-induced obesity and metabolic disorders has been recognized (Bartelt and Heeren, 2014; Crane et al., 2015; Hasegawa et al., 2018; Ikeda et al., 2017; Lowell et al., 1993; Tseng et al., 2010), a comprehensive understanding of developmental origins and regulatory mechanism of beiging is still missing, partially due to the cell-type complexity in adipose tissues. It has been appreciated that beige adipocytes arise by both de novo adipogenesis from progenitor cells and direct conversion/transdifferentiation from white adipocytes (Barbatelli et al., 2010; Shao et al., 2019; Wang et al., 2013). While previous profiling studies have identified some specific markers (for example, CD137 and CD81) for progenitors of beige adipocytes (Oguri et al., 2020; Wu et al., 2012), the cellular origin of beige adipocytes derived from direct conversion remained elusive.

Wnt/β-catenin (canonical) signaling pathway plays a fundamental role in cell proliferation, differentiation, and tissue homeostasis. In the presence of Wnts, “destruction complex” including glycogen synthase kinase 3 (GSK-3) is inhibited and cytoplasmic β-catenin translocates into the nucleus and interacts with TCF/LEF transcription factors, which is the hallmark of the canonical Wnt signaling activation, to activate downstream target genes (Cadigan and Liu, 2006; Dale, 1998). In addition, β-catenin nuclear translocation can also be triggered by other intracellular factors such as AKT and Gα_s_, leading to the activation of β-catenin-mediated signaling in a ligand- and receptor-independent manner (Fang et al., 2007; Regard et al., 2011). Despite the consensus that Wnt/β-catenin signaling imposes negative effects on adipogenesis by inhibiting Pparγ (Longo et al., 2004; Ross et al., 2000; Waki et al., 2007), clues have been pointing to potential roles of Wnt/β-catenin signaling in adipogenesis and adipose tissue function. For instance, β-catenin and TCF7L2, two key effectors of canonical Wnt signaling pathway, are expressed by mature adipocytes. Adipocyte-specific mutations in *Ctnnb1* (encoding β-catenin), *Tcf7l2*, or *Wls* (encoding Wntless) led to impaired adipogenesis (Bagchi et al., 2020; Chen et al., 2020; Chen et al., 2018). Particularly, loss of *Tcf7l2* in mature adipocytes gave rise to adipocyte hypertrophy, inflammation, as well as systemic glucose intolerance and insulin resistance, implying an important biological role for Wnt/β-catenin signaling in adipose function (Chen et al., 2018; Geoghegan et al., 2019). However, direct evidence for active Wnt/β-catenin signaling in adipocytes is lacking.

In the current studies, we have revealed an unexpected adipocyte population that is marked by active Wnt/β-catenin signaling intracellularly, named as Wnt^+^ adipocytes. We found that the Wnt^+^ adipocytes are derived from progenitor lineage that is distinct from that of classical adipocytes and emerge at embryonic stage. Using single-cell transcriptomics and chromatin accessibility profiling assays, we showed that these Wnt^+^ adipocytes distinguish from other conventional ones with respect to molecular and genomic signatures and are highlighted by thermogenic properties. Cold exposure studies uncovered that the Wnt^+^ fat cells play a crucial role in thermogenic response via not only undergoing direct conversion to beige adipocytes but also exerting an indispensable role in beige fat recruitment, representing an important type of thermogenesis-regulatory cells. Finally, gain- and loss-of-function studies further revealed the Wnt^+^ adipocytes as a beneficial metabolic determinant in blood glucose control. The fact that this novel population of adipocytes may also exist in humans implicates it as a potential therapeutic target for metabolic diseases.

## RESULTS

### A unique fraction of murine adipocytes displays Wnt/β-catenin signaling activity

In an unrelated study on the effect of the canonical Wnt signaling on osteogenic differentiation of murine bone marrow stromal cells (BMSCs), we used BMSCs from the well-defined Wnt/β-catenin signaling specific reporter mouse line *TCF/Lef:H2B-GFP* (hereafter *T/L-GFP*) that allows for real-time monitoring of Wnt/β-catenin activity at single-cell resolution (Ferrer-Vaquer et al., 2010). Adipogenic differentiation of *T/L-GFP* BMSCs was conducted in parallel as a negative control for the activation of Wnt/β-catenin signaling. Surprisingly, we observed repeatedly (n > 10) the presence of a small population of GFP-positive cells containing lipid droplets (data not shown). This unexpected observation prompted us to conduct a careful survey for the presence of active Wnt/β-catenin signaling in any developing and mature adipocytes of fat depots from *T/L-GFP* mice. In addition to those tissues and cells known to manifest active Wnt/β-catenin signaling (Ferrer-Vaquer et al., 2010), we virtually observed a subset of Perilipin^+^ adipocytes that exhibited Wnt/β-catenin signaling activity interspersed within various fat depots, including interscapular BAT (iBAT), inguinal WAT (iWAT), epididymal WAT (eWAT), and bone marrow (BM) in adult mice (Figure 1A). Since adipose tissues possess multiple types of non-adipocytes (Cinti, 2005), and the nuclei are often squeezed at the edge of mature white adipocytes, it is difficult to determine the exact hosts of the GFP^+^ nuclei in white adipose tissues. To confirm that these Wnt/β-catenin-positive cells are indeed adipocytes, we created a *Tcf/Lef-CreERT2* transgenic allele in which the *CreERT2* cassette was placed under the control of the same regulatory elements as that used in the *T/L-GFP* allele (Figure 1-figure supplement 1A). This *Tcf/Lef-CreERT2* allele, upon compounding with *Rosa26R*^mTmG^ reporter allele followed by tamoxifen administration, indelibly labeled the membrane of a fraction of adipocytes, as seen in freshly isolated cells and in sectioned iWAT and iBAT (Figure 1-figure supplement 1B, C, E, F). Moreover, addition of the *T/L-GFP* allele to *Tcf/Lef-CreERT2*;*Rosa26R*^mTmG^ mice produced adipocytes that were tagged by both membrane-bound and nucleus-localized GFP (Figure 1-figure supplement 1D). Therefore, we identified a population of adipocytes that displays active Wnt/β-catenin cascade, and accordingly, we referred to these cells as Wnt^+^ adipocytes.

**Figure 1.**
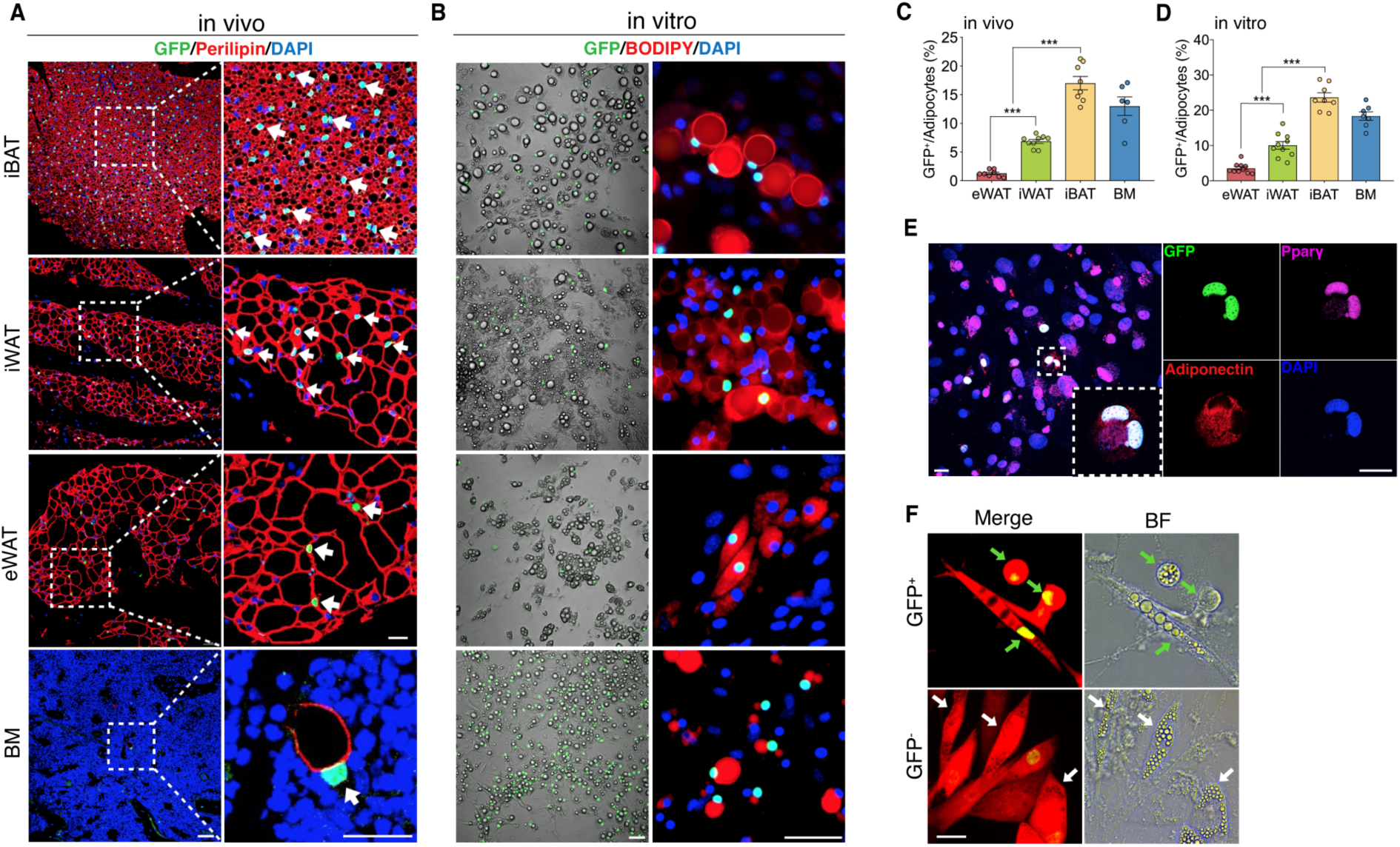
The existence of adipocytes exhibiting active Wnt/β-catenin signaling. (**A** and **B**) Immunofluorescent and microscopy images of active Wnt/β-catenin *in vivo* (**A**) and *in vitro* (**B**), indicated by GFP expression from the *TCF/Lef:H2B-GFP* allele, in adipocytes marked by Perilipin or BODIPY (red). Nuclei were stained with DAPI (blue). Scale bars, 50 μm; close-up scale bars, 20 μm. (**C**) Quantification of Wnt^+^ adipocytes among total adipocytes in various fat depots of adult male mice in (**A**). The levels of mRNA expression are normalized to that of *36B4*. *n* = 6-9 mice. (**D**) Quantification of Wnt^+^ adipocytes among total adipocytes induced from SVF cells derived from adult male fat depots and BM stroma in (**B**). mRNA expression relative to *36B4*. *n* = 7-10 independent experiments. (**E**) Immunofluorescent staining of Pparγ (purple) and Adiponectin (red) in cultured GFP^+^ adipocytes derived from bone marrow in (**B**). *n* = 3 independent experiments. Scale bars, 20 μm. (**F**) Representative images of GFP^+^ (green arrows) and GFP^-^ (white arrows) adipocytes induced from human bone marrow stromal cells infected with *TCF/Lef:H2B-GFP* reporter lentiviral virus. *n* = 2 independent experiments, 3 independent wells each. Scale bar, 20 μm. Data are mean ± s.e.m., \*\*\**P* < 0.001, one-way ANOVA followed by Tukey’s test.

To chart a census of Wnt^+^ adipocytes within distinct mouse fat depots, we quantified Wnt^+^ fat cells in *T/L-GFP* mice. We observed that the adipocytes expressing nuclear GFP count for various percentages of total fat cells in different fat depots, with the highest level (17.02% ± 3.06%) in iBAT, the lowest one (1.28% ± 0.56%) in eWAT, and relatively abundant percentage (6.86% ± 0.98%) in iWAT, a representative beiging site (Nedergaard and Cannon, 2014), of adult male mice (Figure 1C). In addition, Wnt^+^ adipocytes could be detected as early as embryonic day 17.5 (E17.5) (Figure 1-figure supplement 2A, B) and the proportions varied in different fat depots over the course of postnatal stages (Figure 1-figure supplement 2E, F). Interestingly, sex and age appeared to have an impact on the amounts of Wnt^+^ adipocytes, as the percentage of Wnt^+^ adipocyte in female fat depots was around half as compared to their male counterparts and was dramatically reduced in aged male mice (Figure 1-figure supplement 2C-F).

### Wnt^+^ adipocytes can be derived from multiple cellular sources including human BMSCs

To characterize Wnt^+^ adipocytes, we set out to determine if the resident SVF cells within adipose tissues constitute the source of Wnt^+^ adipocytes. Using standard white fat pro-adipogenic induction protocol, we readily induced Wnt^+^ adipocytes *in vitro* from SVFs derived from iBAT, iWAT, eWAT, as well as BM stroma, as confirmed by staining of the general adipocyte markers Pparγ and Adiponectin (Figure 1B and E). The induced Wnt^+^ adipocytes exhibited similar morphology compared to Wnt^-^ (non-GFP labeled) ones and also made up a very similar percentage of total induced fat cells as they are present in their corresponding fat depot (Figure 1D), suggesting that the Wnt^+^ adipocytes we observed *in vivo* are derived from the adipogenic precursors among SVFs. Given the fact that Wnt^+^ adipocytes are present in embryos, we asked if embryonic mesenchymal stem cells can differentiate to Wnt^+^ adipocytes as well. Notably, we could induce E13.5 mouse embryonic fibroblasts (MEFs) to differentiate into Wnt^+^ adipocytes *in vitro* (Figure 1-figure supplement 2G), indicating an early developmental origin of Wnt^+^ adipocyte precursors.

To determine if Wnt^+^ adipocytes could also be induced from human stromal cells, we transfected human primary BMSCs with an mCherry-expressing lentiviral vector carrying a *TCF/Lef:H2B-GFP* reporter cassette (Figure 1-figure supplement 3A) and then cultured these cells with pro-differentiation medium. As the positive control, pre-osteocytes/osteocytes differentiated from infected BMSCs under pro-osteogenic conditions exhibited overlapped mCherry expression and nuclear Wnt/β-catenin signaling activity (Figure 1-figure supplement 3B), validating the effectiveness of the transduction system. Importantly, under the pro-adipogenic medium, we observed GFP-tagged adipocytes induced from the infected BMSCs (Figure 1F), indicating the differentiation of Wnt^+^ adipocytes from human stromal cells. Together, these results suggest that Wnt^+^ fat cells appear to constitute a widespread adipocyte population that originates from embryonic stage and exists in mice and possibly in humans.

### Wnt/β-catenin signaling is activated in Wnt^+^ adipocytes in an intracellular manner

We next addressed how Wnt^+^ adipocytes develop during adipogenesis by conducting real-time monitoring on behaviors of induced Wnt^+^ adipocytes *in vitro*. No visible GFP expression was found in SVFs in culture prior to pro-adipogenic induction (Figure 2A). Nuclear GFP expression was initially detected in differentiated adipocytes (defined by the presence of lipid droplets) after 2-day adipogenic induction and the GFP signal of the cells being constitutively monitored over the course of adipogenesis was sustained once it was activated (Figure 2A), indicating that Wnt/β-catenin signaling does not transiently appear but persists in Wnt^+^ adipocytes. Interestingly, after 7-day differentiation, we observed colonized Wnt^+^ and Wnt^-^ adipocytes, respectively, induced from primary BMSCs of *T/L*-*GFP* mice, suggesting distinct cell lineages of these two different adipocyte populations (Figure 2-figure supplement 1A). This conclusion is supported by the observation that induced Wnt^+^ and Wnt^-^ adipocytes, after separation by fluorescence-activated cell sorting (FACS) and subsequent 4-day in culture, did not convert mutually (Figure 2B). The persistent Wnt/β-catenin signaling within Wnt^+^ adipocyte thus represents an intrinsic cascade activity.

**Figure 2.**
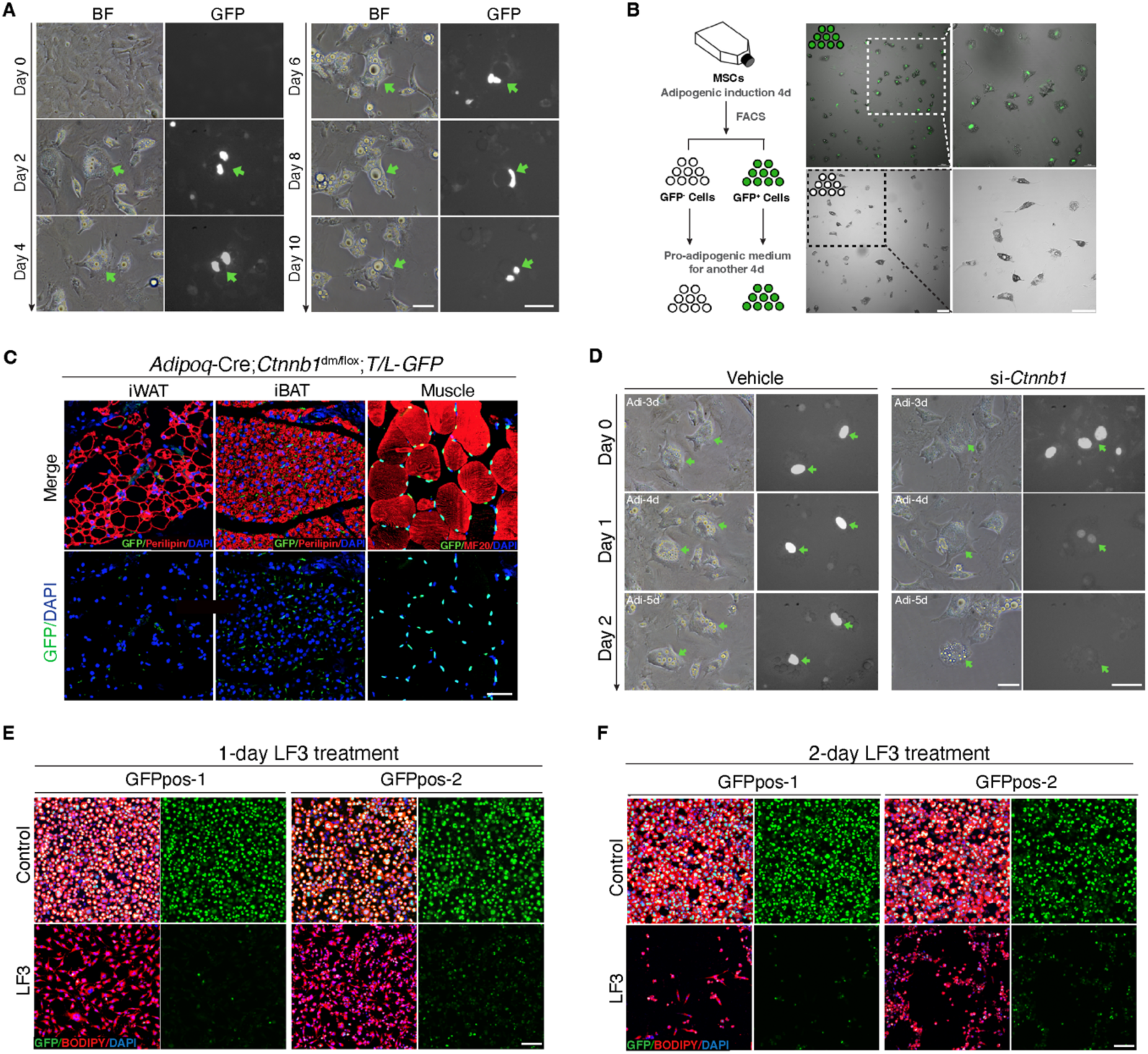
Characterization of mouse Wnt^+^ adipocytes. (**A**) Real-time monitoring microscopy images of Wnt^+^ adipocytes during adipogenesis, which were induced from BMSCs of *T/L-GFP* mice. *n* = 3 independent experiments. Scale bar, 50 μm. (**B**) Schematic of the experiments and microscopy images of separated Wnt^+^ and Wnt^-^ adipocytes by FACS. *n* = 3 independent experiments. Scale bars, 100 μm. (**C**) Immunofluorescent images of iWAT and iBAT of *Adipoq*-Cre;*Ctnnb1*^dm/flox^;*T/L-GFP* mice showing complete absence of Wnt^+^ adipocytes. Muscle cells were included as positive controls for GFP expression. *n* = 3 mice. Scale bar, 50 μm. (**D**) Time-lapse microscopy images of induced Wnt^+^ adipocytes from BMSCs of *T/L*-*GFP* mice with *Ctnnb1* and control siRNA-mediated knockdown. *n* = 2 independent experiments, 3 independent wells each. Scale bars, 50 μm. (**E** and **F**) Immunofluorescent images of Wnt^+^ adipocytes induced from two independent cell lines (GFPpos-1 and -2) with LF3 treatment (50 μM) for one (**E**) and two days (**F**), respectively. LF3 was added into the medium after 3-day pro-adipogenic induction. Note that by 1-day LF3 administration, GFP signals were significantly quenched in Wnt^+^ adipocytes, along with obviously reduced cell number. By 2-day LF3 administration, remarkable cell death of Wnt^+^ adipocytes was seen, compared to controls. Scale bar, 100 μm.

To ensure that the GFP expression in *T/L-GFP* adipocytes both *in vivo* and *in vitro* represents authentic Wnt/β-catenin signaling activity, we first confirmed the presence of active β-catenin and TCF/LEF1 in the nuclei of Wnt^+^ adipocytes within adipose tissues (Figure 2-figure supplement 1B, C). We found that TCF7L2 expression was largely overlapped (∼91%) with Wnt^+^ adipocytes, whereas other TCF proteins (TCF1, TCF3, and LEF1) were barely detected in Wnt^+^ adipocytes (data not shown). To further confirm that the GFP expression by the *T/L-GFP* allele is indeed β-catenin dependent, we first set to abolish β-catenin signaling function *in vivo*. We took advantage of the *Ctnnb1*^dm^ allele that produces truncated β-catenin with abolished transcriptional outputs but retained cell adhesion function (Valenta et al., 2011) and the floxed *Ctnnb1* allele by creating mice carrying adipocyte-specific elimination of β-catenin-mediated signaling. Such mice (*Adipoq*-Cre;*Ctnnb1*^dm/flox^) were crossed to *T/L-GFP* mice, and the adipose tissues of the resultant mice (*Adipoq*-Cre;*Ctnnb1*^dm/flox^;*T/L-GFP*) were subjected to examination of GFP expression by the *T/L-GFP* allele. The results showed complete lack of Wnt^+^ adipocytes in the absence of β-catenin-mediated signaling in both iBAT and iWAT, indicating the absolute requirement of β-catenin-mediated signaling for the GFP expression in Wnt^+^ adipocytes (Figure 2C). Next, we asked if loss of β-catenin could diminish GFP expression in the Wnt^+^ adipocytes by conducting *in vitro* knockdown experiments using short interfering RNA (siRNA) targeting *Ctnnb1* followed by real-time monitoring. *Ctnnb1* knockdown in differentiated Wnt^+^ adipocytes in which GFP expression had been activated virtually quenched the GFP signals, followed by cell shrinkage (Figure 2D). However, surprisingly, we found that DKK1 and IWP-2, both canonical Wnt receptor inhibitors, failed to, but LF3, a molecule that specifically disrupts the interaction between β-catenin and TCF7L2 (Fang et al., 2016), did inhibit GFP expression in adipocytes differentiated from *T/L-GFP* SVFs (Figure 2-figure supplement 1D-G). This result indicates that the transcriptional activity of β-catenin-TCF7L2 complex-mediated intracellular Wnt/β-catenin signaling in Wnt^+^ adipocytes is ligand- and receptor-independent. Remarkably, pharmacological inhibition of Wnt/β-catenin signaling by LF3 also resulted in an overall impaired/delayed adipogenic maturation of SVF-derived adipocytes in a dose-dependent manner (Figure 2-figure supplement 1D). Since most of the SVFs in such culture differentiated into Wnt^-^ adipocytes, this observation suggests the potentially functional importance of Wnt/β-catenin signaling in adipogenesis of Wnt^-^ adipocytes.

To further define the specific roles of the intracellular Wnt/β-catenin signaling in Wnt^+^ adipocytes, we immortalized SVF cells (mBaSVF) derived from iBAT of *T/L-GFP* mouse. By serial limited dilutions, we isolated and established two Wnt^+^ (GFPpos-1 and GFPpos-2) and two Wnt^-^ (GFPneg-1 and GFPneg-2, as controls) adipocyte precursor cell lines from mBaSVF cells, respectively. Of note, LF3 treatment of induced Wnt^+^ adipocytes from both GFPpos-1 and GFPpos-2 cell lines quenched GFP expression in the first day of culture (Figure 2E), followed by massive cell death in the second day (Figure 2F). By contrast, LF3 administration to induced Wnt^-^ adipocytes from precursor cell lines did not affect cell viability but slowed down the maturation of Wnt^-^ adipocytes (Figure 2-figure supplement 2A-D), similar to that seen in SVF induced adipocytes (Figure 2-figure supplement 1D). However, such LF3-treated Wnt^-^ adipocytes, once returned to normal pro-adipogenic medium, resumed full adipogenic capacity compared to controls (Figure 2-figure supplement 2E, F), indicating that LF3 treatment does not impair but delays the maturation of Wnt^-^ adipocytes. As controls, the same dose of LF3 showed no impacts on uninduced Wnt^+^ and Wnt^-^ precursor cell lines (Figure 4-figure supplement 1E).

### Wnt^+^ adipocytes are diverse from conventional ones with distinct molecular and genomic characteristics

To further distinguish Wnt^+^ adipocytes from Wnt^-^ fat cells and explore the global diversity at molecular and genomic levels, we performed single-cell RNA sequencing analysis (scRNA-seq) and single-cell assay for transposase-accessible chromatin sequencing (scATAC-seq) on FACS separated Wnt^+^ and Wnt^-^ adipocytes induced from iWAT-and iBAT-derived SVF cells, respectively (Figure 3A). After filtering for quality and excluding nonadipocytes (negative for *Adiponectin* expression), bioinformatic analyses classified the input Wnt^+^ and Wnt^-^ adipocytes into distinct clusters, indicating two different types of adipocytes (Figure 3B-E, Figure 3-figure supplement 1A-F). Notably, although the mRNA expression of *Ucp1* was undetectable based on our single-cell transcriptomic data, violin plots showed that several thermogenesis-related genes such as *Cox8b* and *Elovl3* were present at significantly higher levels in Wnt^+^ adipocytes that were subject to the white fat differentiation condition without browning stimuli (Figure 3F), indicative of potentially thermogenic character. Moreover, *Cyp2e1*, a molecule identified as a marker gene in thermogenic regulation (Sun et al., 2020b), and *Cidea* were exclusively expressed in a subset of Wnt^+^ adipocytes induced from iWAT- and iBAT-derived SVFs, as confirmed by immunofluorescent staining (Figure 3D-F, Figure 3-figure supplement 1G, H). These facts link Wnt^+^ adipocytes to thermogenic function in adipose tissues. Enrichment pathway analyses of differentially expressed genes (DEGs) also suggests the primary functions of this population of adipocytes in the regulation of adipogenesis, fatty acid metabolism, and mTORC1 signaling (Figure 3G-J). Thus, these results demonstrated that this novel population of intracellular Wnt/β-catenin signaling driven adipocytes is distinct from the classical adipocytes at molecular and genomic levels.

**Figure 3.**
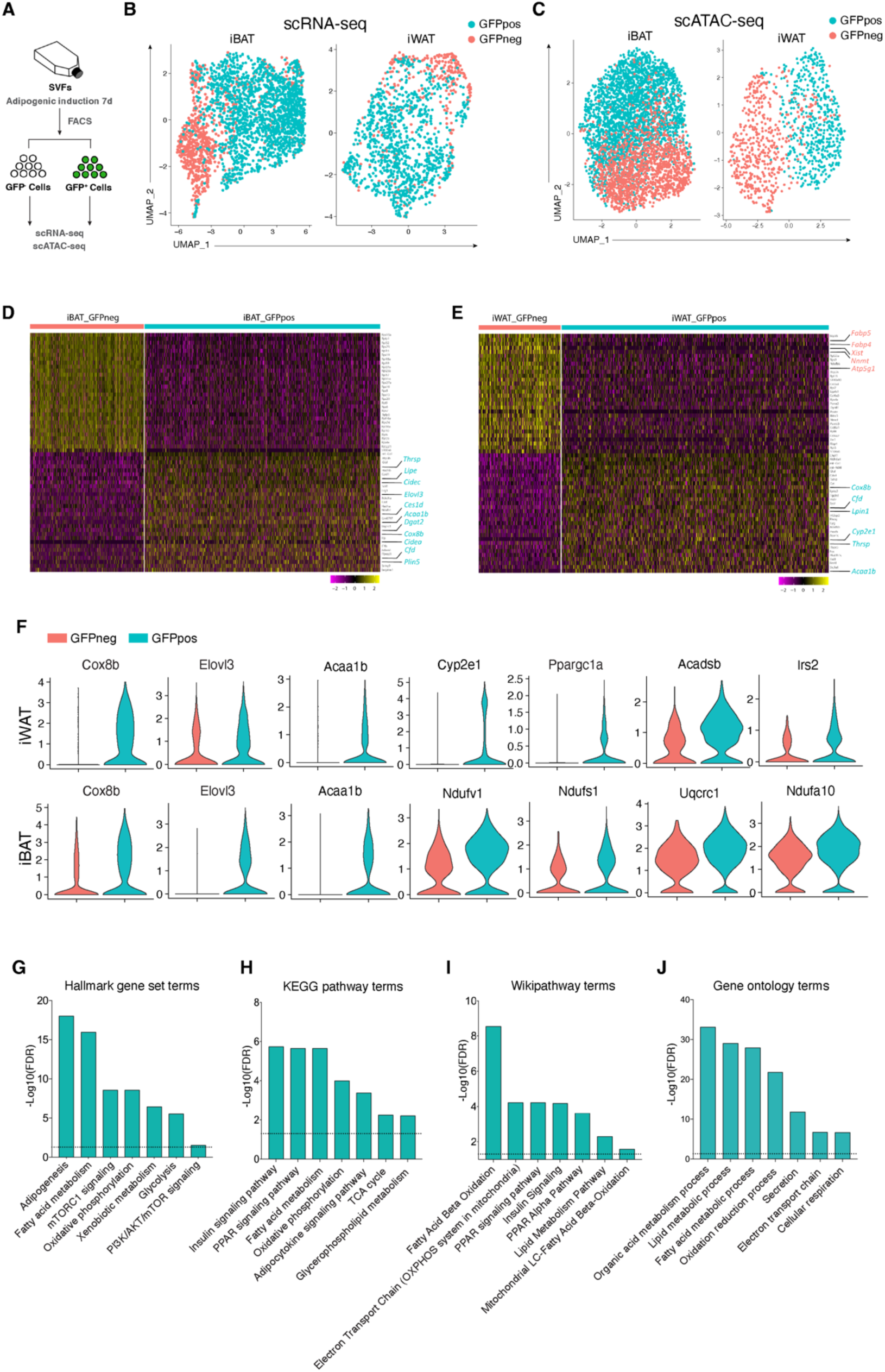
Distinct molecular and genomic signatures of Wnt^+^ and Wnt^-^ adipocytes. (**A**) Schematic of scRNA-seq and scATAC-seq experiments on SVF-induced adipocytes. (**B**) Uniform Manifold Approximation and Projection (UMAP) visualization of 2,537 adipocytes from iBAT (1,710 GFP^+^ and 827 GFP^-^) and 1,345 adipocytes from iWAT (984 GFP^+^ and 361 GFP^-^) in scRNA-seq. (**C**) UMAP visualization of 4,302 adipocytes from iBAT (1,727 GFP^+^ and 2,575 GFP^-^) and 1,383 adipocytes from iWAT (562 GFP^+^ and 821 GFP^-^) in scATAC-seq. (**D** and **E**) Heat maps of expression of top 30 signature genes (Supplementary table 2) in iBAT- (**B**) and iWAT-derived (**C**) Wnt^+^ and Wnt^-^ adipocytes in scRNA-seq. (**F**) Violin plots of induced Wnt^+^ and Wnt^-^ adipocytes showing the distribution of normalized expression values of some representative genes in scRNA-seq. (**G**-**J**) Hallmark gene sets (**G**), KEGG pathway (**H**), Wikipathway (**I**), and GO Biological Processes ontology (**J**) analyses of DEGs enriched in iWAT-derived Wnt^+^ adipocytes in scRNA-seq (Supplementary table 3).

### Insulin/AKT/mTORC1 signaling is required for activation of intracellular Wnt/β-catenin cascade within Wnt^+^ adipocytes

PI3K/AKT/mTOR signaling modulated by insulin is manifested for promoting cytoplasmic β-catenin accumulation through GSK-3β phosphorylation (Hermida et al., 2017; Schakman et al., 2008). This notion upon signaling crosstalk drew our attention because insulin signaling plays pivotal roles in mediating adipogenic differentiation and functionality, and our enrichment pathway analyses also implicate a link to mTORC1 signaling (Figure 3G-J). Accordingly, we first assessed AKT signaling activities in induced Wnt^+^ and Wnt^-^ adipocytes. Immunoblot assay showed that GFPpos-derived Wnt^+^ adipocytes exhibited markedly higher levels of AKT phosphorylation than that from GFPneg-derived Wnt^-^ adipocytes and from mBaSVF-induced adipocytes as unbiased fat cell control (Figure 4A). Phosphorylated GSK-3β and 4E-BP1 (Eukaryotic translation initiation factor 4E-binding protein 1), a known substrate of mTOR signaling pathway (Beretta et al., 1996), were also found preferentially higher in Wnt^+^ adipocytes as compared to controls (Figure 4B). These results demonstrate enhanced AKT/mTORC1 cascade activity and insulin sensitivity in Wnt^+^ fat cells.

**Figure 4.**
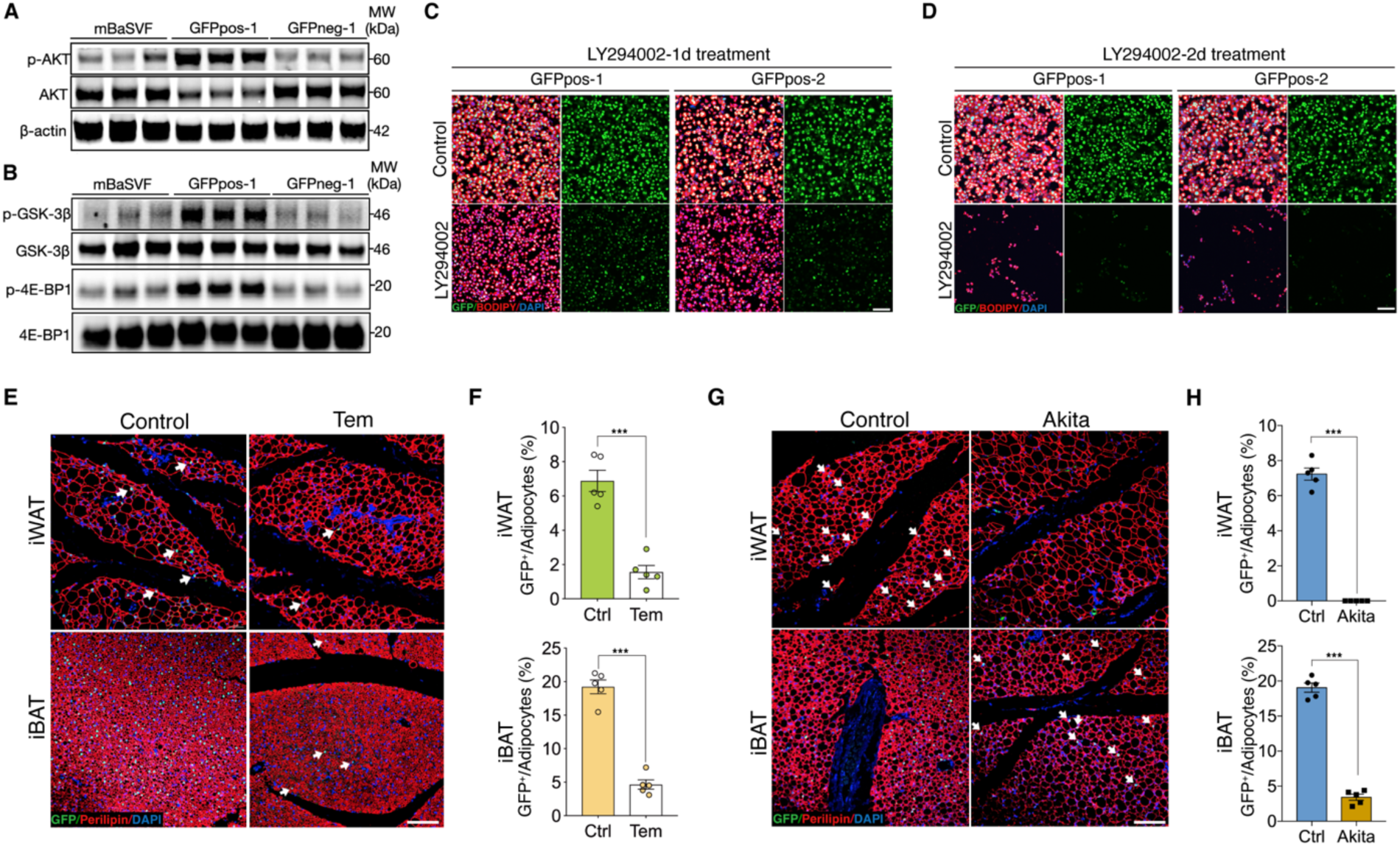
AKT/mTOR cascade is indispensable to trigger intracellular β-catenin signaling in Wnt^+^ adipocytes. (**A**) Western blot analysis showing protein levels of AKT, phosphorylated AKT, and β-actin in adipocytes induced from mBaSVF, GFPpos-1, and GFPneg-1 cell lines. *n* = 3 independent experiments. (**B**) Western blot analysis showing protein levels of GSK-3β, phosphorylated GSK-3β, 4E-BP1, and phosphorylated 4E-BP1 in adipocytes induced from mBaSVF, GFPpos-1, and GFPneg-1 cell lines. *n* = 3 independent experiments. (**C** and **D**) Immunofluorescent images of Wnt^+^ and Wnt^-^ adipocytes induced from two independent precursor cell lines (GFPpos-1 and -2) with LY294002 treatment (14 μM) for one (**C**) and two days (**D**), respectively. LY294002 was added into the medium after 3-day pro-adipogenic induction. LY294002 treatment diminished GFP signals prior to causing marked cell death in Wnt^+^ adipocytes. *n* = 5 independent experiments. Scale bar, 100 μm. (**E**) Immunofluorescent images of iWAT and iBAT of *T/L-GFP* mice treated with or without Temsirolimus (Tem). *n* = 5 mice. Scale bar, 100 μm. (**F**) Quantification of Wnt^+^ adipocytes among total adipocytes in (**E**). (**G**) Immunofluorescent images of iWAT and iBAT of male *Ins2^Akita^*;*T/L-GFP* mice at 8 weeks of age. *n* = 5 mice. Scale bar, 100 μm. (**H**) Quantification of Wnt^+^ adipocytes among total adipocytes in (**G**). Data are mean ± s.e.m., \*\*\**P* < 0.001, unpaired Student’s t-test.

To test if AKT/mTOR signaling is responsible for intracellular β-catenin-mediated signaling activation in Wnt^+^ adipocytes, we started with treating iBAT-derived SVF cells with LY294002, a selective PI3K signaling inhibitor, under pro-adipogenic conditions. Again, dramatically reduced number of Wnt^+^ adipocytes and blunted adipogenesis were observed in a dose-dependent manner (Figure 4-figure supplement 1A, B). Consistently, LY294002 administration to GFPpos-induced Wnt^+^ adipocytes eliminated GFP expression and caused substantial cell death subsequently but did not impact on the survival of GFPneg-derived adipocytes (Figure 4C and D, Figure 4-figure supplement 1C-E). To determine a causal role of mTOR signaling in the intracellular activation of β-catenin-mediated signaling in adipocytes *in vivo*, we subjected *T/L-GFP* adult mice to an mTOR-specific inhibitor Temsirolimus treatment, which dramatically reduced Wnt^+^ adipocyte number in both iWAT and iBAT compared to vehicle-injected controls (Figure 4E and F). These results demonstrate the dependence of the intracellular β-catenin signaling activity in adipocytes on AKT/mTOR signaling, which is required for cell survival.

To further validate the requirement of insulin signaling in Wnt^+^ adipocyte development *in vivo*, we employed *Ins2^Akita^* mice (the so-called Akita mice) in which insulin secretion is profoundly impaired due to genetic defect in the insulin 2 gene (Yoshioka et al., 1997) by crossing them to *T/L-GFP* mice. As expected, Wnt^+^ adipocytes were completely absent in iWAT and substantially reduced in number in iBAT (3.34% ± 0.55%) of 8-week-old male Akita mice (Figure 4G and H), demonstrating an indispensable effect of insulin-induced AKT/mTOR signaling on Wnt^+^ adipocytes differentiation and/or maintenance.

### Wnt^+^ adipocytes are required for initiating adaptive thermogenesis in both cell autonomous and non-cell autonomous manners

The potentially highly metabolic and thermogenic characters in Wnt^+^ adipocytes prompted us to investigate the adaptive thermogenic role of Wnt^+^ adipocytes. We started with the examination of mitochondrial activities in SVF-derived adipocytes *in vitro* and found that Wnt^+^ adipocytes, along with those closely adjacent Wnt^-^ ones, exhibited pronounced lower levels of mitochondrial membrane potential, indicative of higher uncoupling rate, as compared to those Wnt^-^ fat cells located relatively away from Wnt^+^ adipocytes (Figure 5A and B). In addition, significantly higher levels of oxygen consumption rate (OCR) were detected in GFPpos-induced Wnt^+^ adipocytes as compared to those GFPneg-derived Wnt^-^ adipocytes and mBaSVF-induced fat cells (Figure 5C, Figure 5-figure supplement 1A, B), further demonstrating a higher mitochondrial respiration capacity of Wnt^+^ adipocytes.

**Figure 5.**
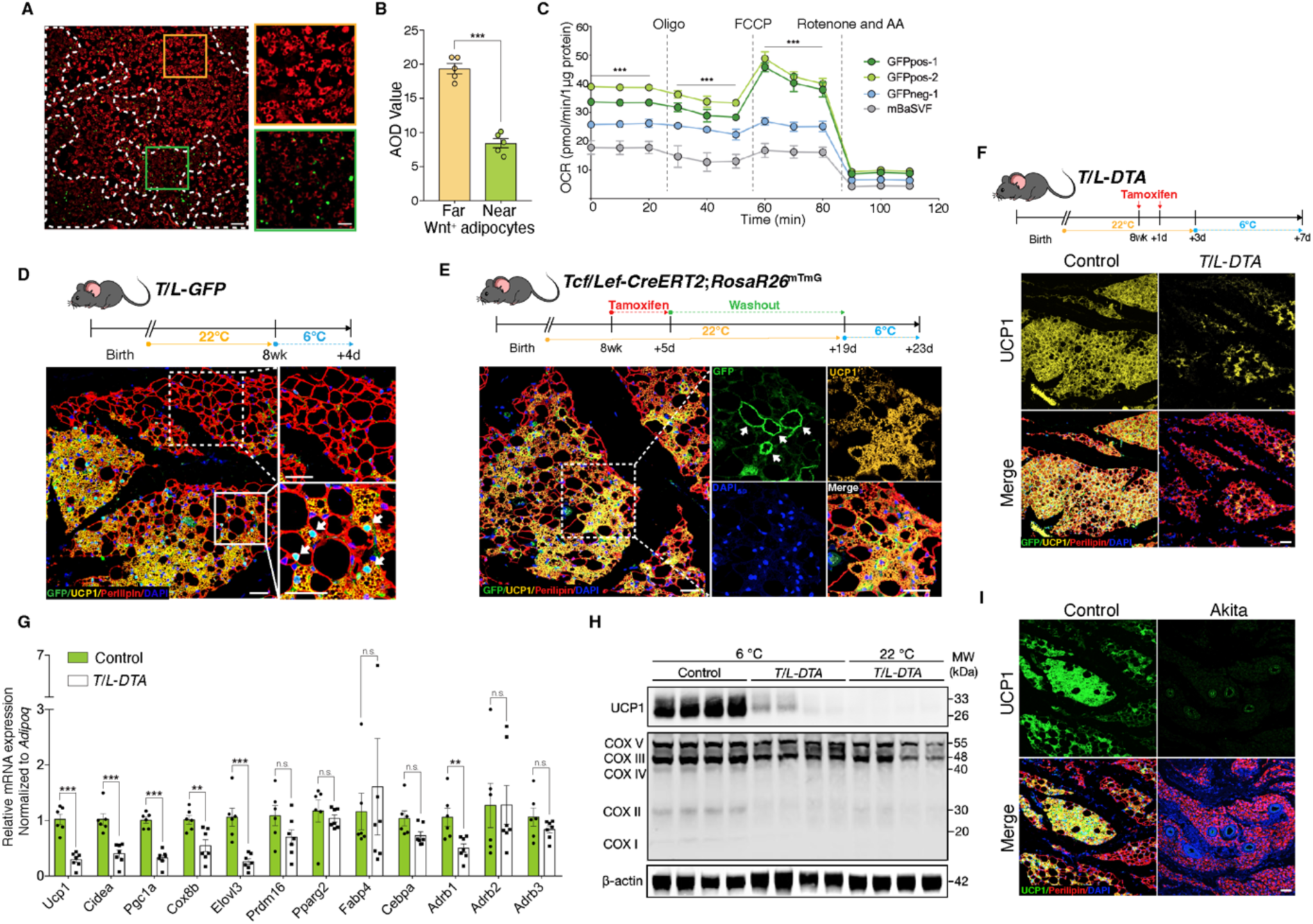
Wnt^+^ adipocytes are essential for initiating adaptive thermogenesis. (**A**) Immunofluorescent staining of mitochondrial membrane potentials in Wnt^+^ adipocytes induced from iBAT-derived SVF cells of *T/L-GFP* mice. *n* = 5 independent experiments. Scale bar, 100 μm; close-up scale bar, 20 μm. (**B**) Quantification of staining in (**A**). AOD, average optical density. Data are mean ± s.e.m., unpaired Student’s *t*-test; \*\*\**P* < 0.001. (**C**) OCR plots of four groups of adipocytes differentiated from GFPpos-1, GFPpos-2, GFPneg-1, and mBaSVF cell lines, respectively. *n* = 3 independent experiments. (**D**) Immunofluorescent staining of iWAT from *T/L*-*GFP* mice with 4-day thermal challenge showing close association of Wnt^+^ adipocytes with UCP1^+^ beige adipocytes. *n* = 5 mice. Scale bar, 100 μm. (**E**) Immunofluorescent staining of iWAT from tamoxifen-treated *Tcf/Lef*-*CreERT2*;*Rosa26R*^mTmG^ mice after 4-day cold exposure. *n* = 8 mice. Scale bar, 50 μm. (**F**) Immunofluorescent staining of iWAT from tamoxifen-treated *T/L*-*DTA* mice after 4-day cold exposure. Before cold challenge, mice were rested for 48 hours after final tamoxifen treatment. *n* = 7 mice. Scale bar, 50 μm. (**G**) Quantitative RT-PCR analysis of gene expression in iWAT from control (*Fabp4-Flex-DTA*) and *T/L*-*DTA* mice in (F). *n* = 6 and 7 mice. Levels of mRNA expression are normalized to that of *Adipoq*. (**H**) Western blot analysis showing protein levels of UCP1, OXPHOS complexes, and β-actin in iWAT from tamoxifen-treated controls and *T/L*-*DTA* mice under cold (6°C) and ambient (22°C) temperatures. *n* = 4 mice. (**I**) Immunofluorescent staining for UCP1 (green) and Perilipin (red) in the iWAT from control and Akita mice. Scale bars, 50 μm. Data are mean ± s.e.m., \**P* < 0.05, ** *P* < 0.01, \*\*\**P* < 0.001, n.s., not significant, two-way repeated ANOVA followed by Bonferroni’s test (**C**) or unpaired Student’s t-test (**G**).

To explore the possible involvement of Wnt^+^ adipocytes in adaptive thermogenesis *in vivo*, adult *T/L-GFP* male mice were subjected to 6°C temperature for 2 days to initiate beiging. After cold exposure, we observed the presence of a subset of UCP1-expressing beige adipocytes that were also GFP-positive (Figure 5-figure supplement 1C), demonstrating that a portion of beige fat cells arises from Wnt^+^ adipocyte lineage. Similar results were seen in mice treated with β3-adrenergic receptor agonist CL316,243 (Figure 5-figure supplement 1C), despite the unaltered proportion of Wnt^+^ adipocytes in iWAT in response to cold stress or β3-adrenergic receptor agonist treatments (Figure 5-figure supplement 1D). Remarkably, as cold exposure was prolonged to 4 days, the number of Wnt^-^ beige adipocytes increased dramatically, but the majority of, if not all, UCP1-labeled Wnt^-^ beige adipocytes were found present neighboring to Wnt^+^ adipocytes (Figure 5D). UCP1^+^/Wnt^-^ beige adipocytes were rarely seen away from Wnt^+^ adipocytes within iWAT, implicating a regulatory role of Wnt^+^ adipocytes in beige fat biogenesis. Together with the results from mitochondrial activity assays, these observations suggest a paracrine function of Wnt^+^ adipocytes in modulating beige fat formation.

To determine whether those Wnt^+^ beige adipocytes are converted directly from Wnt^+^ adipocytes under cold conditions, we conducted lineage-tracing studies using *Tcf/Lef-CreERT2*;*Rosa26R*^mTmG^ mice. After pre-treatment with tamoxifen and a 2-week washout at ambient temperature, *Tcf/Lef-CreERT2*;*Rosa26R*^mTmG^ mice were subjected to cold challenge for 4 days. These mice manifested the presence of mGFP and UCP1 co-labeled adipocytes in iWAT (Figure 5E), providing unambiguous evidence for the cell autonomous contribution of Wnt^+^ adipocytes to beiging.

To establish a non-cell autonomous role for Wnt^+^ adipocytes in beige fat biogenesis, we generated a *Fabp4-Flex-DTA* mouse model (Figure 5-figure supplement 1E), a diphtheria-toxin-induced depletion system that allows targeted ablation of Wnt^+^ adipocytes upon crossing to *Tcf/Lef-CreERT2* mice (*T/L-DTA* mice) and tamoxifen administration, which eliminated about 87% of Wnt^+^ adipocytes in iWAT (Figure 5-figure supplement 1F). Tamoxifen-treated *T/L-DTA* mice appeared normal and did not exhibit lipodystrophic phenotype by histological examination under regular chow diet (data not shown). We then challenged Wnt^+^ adipocyte-ablated mice and littermate controls (*Fabp4-Flex-DTA* mice) at 6°C temperature. We first measured the core body temperature of each mouse and found that about 31% *T/L-DTA* mice (5 of 16 mice) developed hypothermia (below 34.5°C) within 60 hours under cold conditions, with the rest of the *T/L-DTA* mice exhibiting comparable core body temperature as the controls (Supplementary table 4). We next examined if the formation of cold-induced beige fat in iWAT of *T/L-DTA* mice would be blunted. After 4-day cold stress, DTA-induced ablation of Wnt^+^ adipocytes led to substantially reduced UCP1-expressing beige adipocytes compared to control littermates (Figure 5F). This markedly attenuated UCP1 expression persisted even after 2 weeks of cold adaptation (Figure 5-figure supplement 1G). The compromised beiging process in iWAT was verified by remarkably decreased expression levels of thermogenic genes assessed by qRT-PCR and western blotting (Figure 5G and H). Since the mRNA levels of the pan-adipogenic marker *Adipoq* (*Adiponectin*) appeared similarly between the genotypes (Figure 5-figure supplement 1H), it was used as internal reference. As shown in Figure 5G, loss of Wnt^+^ adipocytes in *T/L-DTA* mice caused significantly suppressed expression of thermogenesis-related genes specifically but not those general adipogenic markers in iWAT. Because Akita mice lack Wnt^+^ adipocytes in iWAT (Figure 4G and H), similar to *T/L-DTA* ones, we tested whether Akita mice also display an impaired adaptive thermogenic capacity by challenging Akita mice at 6°C. Immunofluorescence staining further validated that cold-induced UCP1^+^ cells were largely absent in iWAT of Akita mice (Figure 5I). Collectively, these results demonstrate an essential role of Wnt^+^ adipocytes in initiating beige adipogenesis under cold conditions via non-cell autonomous effect.

### Wnt^+^ adipocytes enhance systemic glucose handling in mice

Given that the presence of Wnt^+^ adipocyte is highly dependent on insulin signaling, we set to evaluate the potential physiological impact of Wnt^+^ adipocytes on whole-body glucose homeostasis. We started with loss-of-function studies by using *T/L-DTA* mice, which, under regular chow diet, were subjected to glucose tolerance assays 24 hrs after tamoxifen administration. The results showed clearly that *T/L-DTA* mice displayed impaired glucose handling competence as compared to controls despite similar basal glucose levels (Figure 6A), indicating a requirement for Wnt^+^ adipocytes in handing systemic glucose. We next took gain-of-function approach to test if Wnt^+^ adipocytes are sufficient to enhance systemic glucose utilization. For this purpose, immortalized GFPpos and mBaSVF cells were induced to become adipogenic commitment after 2-day in pro-adipogenic medium. Committed cells were mixed with Matrigel and implanted into the left abdomen subcutaneous layer of wild-type C57BL/6J mice under regular chow diet (Figure 6B). These implanted cells differentiated into fully mature adipocytes readily and formed vascularized adipose tissues *in vivo* within 2-week (Figure 6C and D). Glucose tolerance assays were performed on mice 2-week after cell implantation. The results showed that mice bearing implanted Wnt^+^ adipocytes exhibited a significantly enhanced glucose tolerance as compared to mice receiving mBaSVF-derived adipocytes (Figure 6E). Taken together, these results provide appealing evidence for the beneficial impact of Wnt^+^ adipocytes in systemic glucose homeostasis.

**Figure 6.**
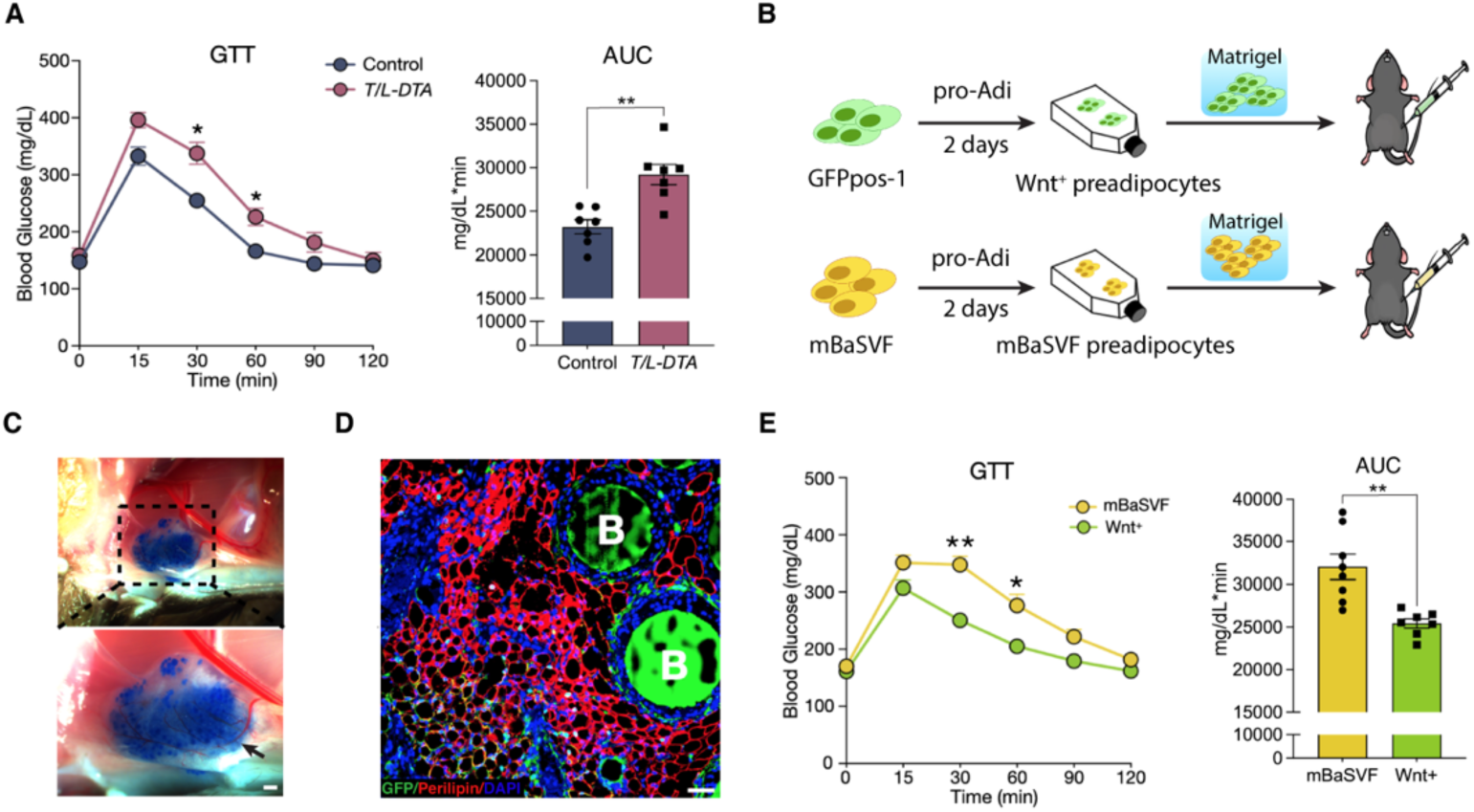
Wnt^+^ adipocytes enhance systemic glucose homeostasis. (**A**) Glucose tolerance test (GTT) with calculated area under the curve (AUC) in tamoxifen-treated control (*Fabp4-Flex-DTA*) and *T/L-DTA* mice on regular chow diet. *n* = 7 mice each. (**B**) Schematic of Wnt^+^ adipocyte gain-of-function studies by cell implantation. (**C**) Photographs of fat pad formed by implanted cells. Blue agarose beads were included to locate the Matrigel pad. Black arrow shows benign vascularization of fat pad within two weeks. Scale bar, 50 μm. (**D**) Immunofluorescent staining for Perilipin showing mature adipocytes and accompanied agarose beads (marked by B) in the ectopically formed fat pad in (**C**). Scale bar, 50 μm. (**E**) GTT with calculated AUC in mice that received implantation of committed pre-adipocytes/adipocytes from mBaSVF (*n* = 8 mice) or GFPpos-1 (*n* = 7 mice) cell lines for two weeks. Data are mean ± s.e.m., \**P* < 0.05, \*\**P* < 0.01, two-way repeated ANOVA followed by Bonferroni’s test. AUC was analyzed by two-tailed *t*-test, *P* = 0.0012 (**A**), 0.0017 (**E**).

## DISCUSSION

Over the past few decades, numerous studies support the notion that the canonical Wnt signaling pathway functions as a powerful suppressor of adipogenesis (Longo et al., 2004; Ross et al., 2000; Waki et al., 2007). Our current studies expand the conventional dogma conceptually by presenting direct evidence for the existence of a population of adipocytes marked by active intracellular Wnt/β-catenin signaling, revealing the diversity of adipocytes. Despite of a previous study reporting that the canonical Wnt signaling is operative persistently in overall mature adipocytes and plays a role in *de novo* lipogenesis (Bagchi et al., 2020), the activity of such canonical Wnt signaling is apparently below the threshold of the *T/L-GFP* sensitivity in those so-called Wnt^-^ adipocytes. However, we demonstrated that the β-catenin-mediated signaling in the Wnt^+^ adipocytes is independent on Wnt ligands and receptors, but is activated intracellularly via signaling crosstalk, in this case, AKT/mTOR signaling. Since β-catenin nuclear translocation and binding to Tcf/Lef1 is regarded as hallmark of canonical Wnt signaling, we still refer this population of fat cells as Wnt^+^ adipocytes. Although if AKT/mTOR signaling cascade can activate Wnt/β-catenin signaling remains contradictory (Baryawno et al., 2010; Cabrae et al., 2020; Ng et al., 2009; Palsgaard et al., 2012; Prossomariti et al., 2020; Zhang et al., 2013), it appears to be cell-type dependent. This population of Wnt^+^ adipocytes was previously neglected, presumably because it is relatively few within fat depots *in vivo* and most of the prior studies were performed in aggregates of mixed cell types through Wnt ligand- and receptor-dependent manners. Importantly, we showed that the intracellularly AKT/mTOR cascade-dependent β-catenin signaling is required for the survival of Wnt^+^ adipocytes *in vitro*. Such intracellular signaling cascade appears to exert distinct function as compared to the ligand-dependent canonical Wnt signaling that regulates lipogenesis in other mature adipocytes (Bagchi *et al*., 2020).

Our single-cell sequencing data showed that Wnt^+^ adipocytes, as compared to Wnt^-^ adipocytes induced from SVF cells of iWAT and iBAT origins, exhibit distinct signatures at molecular and genomic levels, further supporting the Wnt^+^ adipocytes as a unique subpopulation of fat cells in mammals. Because FACS-based selection of mature adipocytes presents a huge challenge and the lipid droplets of cultured adipocytes are much smaller, we had to perform sequencing on *in vitro* cultured cells that were not under their true physiological status. We have attempted to conduct single nucleus RNA sequencing (snRNA-seq) on mature adipocytes isolated from adipose tissues. However, unlike the previous successful studies in which adipocyte nuclei were isolated from adipose tissues simply by one-step enzyme digestion prior to snRNA-seq (Rajbhandari et al., 2019; Sarvari et al., 2020; Sun et al., 2020b), we were not able to obtain desired results due to nuclear membrane breakage after enzyme digestion and following FACS separation of Wnt^+^ and Wnt^-^ adipocyte nuclei. Nevertheless, under the standard white fat pro-adipogenic induction, SVF-induced Wnt^+^ adipocytes, but not Wnt^-^ adipocytes, express several brown/beige fat-selective genes, mirroring the thermogenic feature. In addition, pathway analyses indicate highly enriched insulin signaling pathway in Wnt^+^ adipocytes, as compared to Wnt^-^ fat cells, consistent with the intracellular activation of β-catenin signaling by AKT/mTOR signaling cascade. While many subpopulations of heterogeneous pre-adipocytes/adipocytes have been identified, the classification of each subset replies mainly on the expression of specific genes such as cell surface markers (Hepler et al., 2018; Oguri et al., 2020; Schwalie et al., 2018; Wu et al., 2012). Our current studies, based on the stable feature of an intrinsic cellular signaling identity, provide a new revenue for taxonomy of adipocytes.

In line with their thermogenic feature, as revealed by gene expression profiling, our studies have further uncovered a key role of Wnt^+^ adipocytes in adaptive thermogenesis of adipose tissues. First, Wnt^+^ adipocytes represent at least one of the cell populations that can convert into beige adipocytes directly following cold stress. Interestingly, a conversion of mature white adipocyte to reprogrammed beige characteristics has been postulated to occur predominantly in the “dormant” or “latent” beige fat (i.e., thermogenic inactive state of UCP1 lineage adipocytes within WAT depots), which was thought to form initially through *de novo* beige adipogenesis from progenitors during postnatal development (Paulo and Wang, 2019; Shao et al., 2019). Because UCP1 expression within iWAT starts at postnatal day 14 independent of environmental temperature (Wu et al., 2020), dormant beige adipocytes appear much later in development than Wnt^+^ adipocytes that can be found in embryonic stage. Our recent lineage tracing studies also revealed non-overlapped Wnt^+^ adipocytes and UCP1^+^ dormant beige cells in iWAT of 4-week-old mice (unpublished data). Therefore, together with the observations that Wnt^+^ and Wnt^-^ adipocytes do not convert mutually in cell culture, our findings demonstrate that Wnt^+^ adipocytes differ from dormant beige cells and represent a population of adipocytes that retain the cellular plasticity and are capable of converting to beige adipocytes directly. Second, Wnt^+^ adipocytes do not just simply function in cell autonomous manner in beiging response, but also exert paracrine effect on and are required for beige fat recruitment. Upon cold exposure, UCP1^+^/Wnt^-^ beige fat emerges largely surrounding Wnt^+^ adipocytes, implicating that Wnt^+^ adipocytes serve as a beiging initiator in a paracrine manner. Remarkably, using a targeted cell ablation system, we showed that depletion of Wnt^+^ fat cells in mice leads to severely impaired beige biogenesis in iWAT in response to cold stress. Similarly impaired beiging response was also seen in iWAT of Akita mice that lack Wnt^+^ adipocytes. These observations underscore the functional importance of Wnt^+^ adipocytes in triggering adaptive thermogenic program of adipose tissue. Despite that the critical factors produced by Wnt^+^ adipocytes to trigger beige adipogenesis remain completely unknown, we conclude that Wnt^+^ adipocytes act as a key initiator in beige biogenesis through both cell autonomous (direct conversion) and non-cell autonomous (recruitment) manners, which is thus far only seen in such adipocyte population. However, it is worth noting that even Wnt^+^ adipocytes exhibit a thermogenic character, it does not necessarily mean that the intracellular β-catenin signaling play a direct role in regulating thermogenic function in adipocytes.

The significant contributions of beige adipocytes to improving whole-body metabolism such as glucose homeostasis and countering diet-induced obesity have been well documented (Bartelt and Heeren, 2014; Crane et al., 2015; Harms and Seale, 2013). GWAS has also linked genetic variations in *TCF7L2*, a key component of canonical Wnt pathway, to type 2 diabetes in humans (Chen et al., 2018; Voight et al., 2010). We thus speculated that Wnt^+^ adipocytes, driven by insulin/AKT signaling and functioning a key regulator of beiging in mice, represent a population of beneficial adipocytes and hold the therapeutic potential for metabolic diseases. Accordingly, our results demonstrate whole-body glucose intolerance in mice with targeted ablation of Wnt^+^ adipocytes but notably enhanced glucose utilization in mice receiving implantation of exogenously induced Wnt^+^ adipocytes, highlighting Wnt^+^ adipocytes as a therapeutically appealing candidate of cell source to restore systemic glucose homeostasis. The evidence that Wnt^+^ adipocytes could be induced from human primary BMSCs opens a door for future translational research.

## MATERIALS AND METHODS

### Mice

Animal studies were performed according to procedures approved by the Institutional Animal Care and Use Committee of Tulane University. C57BL/6J wild-type mice (Stock No. 000664), *TCF/Lef:H2B-GFP* mice (Stock No. 013752), *Rosa26R*^mTmG^ mice (Stock No. 007676), and *Ins2^Akita^* mice (Stock No. 003548) were purchased from the Jackson Laboratory and maintained on C57/BL6 background. Mice were raised on a standard rodent chow diet and housed at ambient temperature (22°C) under a 12-hrs light cycles, with free access to food and water. For all experiments, male mice at 8-10 weeks of age were used, unless otherwise stated.

To generate *Tcf/Lef*-*CreERT2* transgenic mice, the coding sequence of the *CreERT2* was cloned into *pTCF/Lef*:*H2B-GFP* vector by replacing *H2B-GFP* sequences under the control of the *hsp68* minimal promoter. *Fabp4*-*Flex*-*DTA* transgenic construct was generated by inserting the coding sequence of diphtheria toxin A (DTA) into the pBS *Fabp4* promoter (5.4kb) polyA vector (Addgene, 11424), flanked by flip-excision (FLEX) switch. Pronuclear injection and embryo transfer were performed following standard protocols at Tulane Transgenic Animal Facility.

*Adipoq*-Cre;*Ctnnb1*^dm/flox^;*T/L-GFP* mice were generated by compounding *Adipoq*-Cre (Stock No. 028020; Jackson Laboratory) allele with *Ctnnb1*^flox^ allele (Stock No. 004152 Jackson Laboratory), *Ctnnb1*^dm^ allele (gift from Dr. K. Basler of the University of Zurich), and *TCF/Lef:H2B-GFP* (*T/L-GPF*) allele. Adult male mice were subjected to adipose tissue harvest for examination of Wnt^+^ adipocytes.

For cold-exposure experiments, mice were singly caged and exposed to cold temperature at 6°C for 2, 4, or 14 consecutive days. For β3-adrenoceptor agonist treatment, adult male mice were injected intraperitoneally with CL316,243 (Sigma-Aldrich, C5976) at 1 mg/kg body weight daily for 4 consecutive days prior to sample collection. Age-matched male littermates were treated intraperitoneally with saline as vehicle controls. For mTOR specific inhibitor administration, adult male mice were injected intraperitoneally with Temsirolimus (Sigma-Aldrich, PZ0020) at 600 μg/kg (dissolved in 40% ethanol) body weight daily for 5 consecutive days prior to sample harvest. Age-matched male littermates were treated intraperitoneally with 40% ethanol as vehicle controls.

To conduct *in vivo* lineage-tracing, *Tcf/Lef*-*CreERT2* allele was compounded with *Rosa26R*^mTmG^ allele to generate *Tcf/Lef*-*CreERT2*;*Rosa26R*^mTmG^ mice that received tamoxifen (dissolved in corn oil) administration intraperitoneally at the dose of 100 mg/kg body weight for 5 consecutive days, and samples were harvested at day 6. For cold exposure study, tamoxifen administrated *Tcf/Lef*-*CreERT2*;*Rosa26R*^mTmG^ mice and littermate controls (*Tcf/Lef*-*CreERT2* mice that received identical tamoxifen treatment) were rested for tamoxifen washout for 2 weeks before they were housed at 6°C for 4 consecutive days.

To ablate Wnt^+^ adipocytes *in vivo*, *Fabp4*-*Flex*-*DTA* mice were crossed with *Tcf/Lef*-*CreERT2* mice to generate *Tcf/Lef*-*CreERT2*;*Fabp4*-*Flex*-*DTA* mice (*T/L-DTA* mice), which received 2-day tamoxifen administration via intraperitoneal injection at the dose of 150 mg/kg body weight. This dose of 2-day tamoxifen administration was found to ablate significantly ablate Wnt^+^ adipocytes (about 87%) in iWAT of *T/L-DTA* mice (Figure 5-figure supplement 1F). Forty-eight hours later, *T/L-DTA* mice and tamoxifen-treated littermate controls (*Fabp4*-*Flex*-*DTA* mice) were housed at 6°C for 2 days, respectively.

### Isolation of mouse adipose stromal vascular fractions (SVFs) and bone marrow stromal cells (BMSCs)

All cells were isolated from adult *TCF/Lef:H2B-GFP* male mice, unless otherwise specified. For the isolation of adipose SVF cells, fat tissues were dissected into ice-cold PBS and minced with scissors in a sterile 5 ml tube containing 4 ml of either iBAT digestion buffer (1X HBSS, 3.5% BSA, and 2 mg/ml collagenase type II) and incubated at 37°C for 1 hr under agitation or WAT digestion buffer (1X DPBS, 1% BSA, 2.5 mg/ml dispase and 4 mg/ml collagenase D) for 40 min. The digestion mixture was passed through a 100 µm cell strainer into a 50 ml tube. Digestion was stopped by adding 15 ml PBS containing 2% fetal bovine serum (FBS; Gibco, 10270106) and centrifuged at 500 g for 5 min at room temperature. The supernatant was aspirated, and red blood cells were lysed by incubating the SVF pellet with 2 ml RBC lysis buffer (Invitrogen, 00433357). The number and viability of cells in the suspension were determined using Trypan Blue stain according to the manufacturer’s recommendations.

For isolation of BMSCs, femurs and tibias were carefully disassociated from muscle, ligaments, and tendons on ice, and then transferred into a sterile 5 ml tube with DPBS containing 2% FBS. Both ends of the bones were cut to expose the interior of the marrow shaft, and bone marrow was flushed out with MesenCult^TM^ Expansion medium (STEMCELL, 05513) using a 6 ml syringe and a #23 gauge needle and collected into a 50 ml tube. Cells were gently resuspended and filtered through a 100 mm filter into a collection tube and cell number and viability were determined as described above.

### Fluorescence-activated cell sorting (FACS)

Freshly isolated single-cell suspension of SVFs was diluted to 1 x 10^7^ cells/ml with FASC sorting buffer (DPBS with 2% FBS, 1 mM EDTA) and the following fluorophore-conjugated antibodies were added: CD31/PE, CD45/PE, and Ter119/PE to enrich lineage^-^ (Lin^-^) populations. Cells were incubated with a cocktail of antibodies on ice for 30 min protected from light and subjected to FACS using a Sony SH800 cell sorter. The cells were selected based on their size and complexity (side and back scatter), and then subjected to doublet discrimination to obtain single-cell signal. The Lin^-^ (CD31^-^CD45^-^TER119^-^) population was sorted out and plated at the density of 4 x 10^3^ cells/cm^2^ for further pro-adipogenic induction.

To sort out Wnt^+^ (GFP^+^) and Wnt^-^ (GFP^-^) adipocytes differentiated from the SVFs for scRNA-seq and scATAC-seq, 7-AAD was used for viable cell gating. Flow cytometric sorting experiments for adipocytes were performed using FACS-Aria II flow cytometer (BD Biosciences).

### Development of immortalized precursors of Wnt^+^ and Wnt^-^ adipocytes

To establish clonal cell lines of immortalized precursors of Wnt^+^ and Wnt^-^ adipocytes, we generated a lentiviral shuttle vector *pUltra-hot-LT* that expresses mCherry and Simian Virus 40 Large T antigen (SV40-LT) simultaneously. SV40-LT fragment was amplified from template plasmid pBABE-neo-LargeTcDNA (Addgene, 1780) with primers 5’-gactcatctagagataaagttttaaacagagaggaatctttgcagc -3’ and 5’-gcatacggatcctgtttcaggttcagggggagg - 3’, and the 2145bp PCR product was subsequently digested with XbaI and BamHI. The vector template pUltra-hot (Addgene, 24130) was linearized with XbaI and BamHI as well. Both the digested PCR product and the linearized pUltra-hot vector were gel-purified and ligated with T4 DNA ligase (Thermo Fisher Scientific, EL0011) according to manufacturer’s instructions. The ligated vector is termed *pUltra-hot-LT* as the final shuttle vector.

For lentivirus (LV) production, 60% confluent monolayers of 293-T cells were transfected with LV shuttle vector *pUltra-hot-LT* and the packaging plasmids psPAX2 (Addgene, 12260) and pMD2.G (Addgene, 12259) at a molar ratio of 4:3:1. The 293-T cells were cultured in high-glucose Dulbecco’s modified Eagle’s medium (DMEM; ThermoFisher Scientific, 12430112) with 10% FBS (ThermoFisher Scientific, 16000069), supplemented with 1% Penicillin-Streptomycin (ThermoFisher Scientific, 15140148) and 1% non-essential amino acids (ThermoFisher Scientific, 11140050). The transfections were performed using Helix-IN transfection kit (OZ Biosciences, HX10100) following manufacturer’s instructions. Forty-eight hrs after transfection, the 293-T culture supernatants were harvested and passed through a 0.45-µm pore-sized, 25-mm diameter polyethersulfone syringe filters (Whatman, 6780-2504). LV particles were concentrated from the filtered supernatants using a Lenti-X concentrator kit (Takara, 631232) following manufacturer’s protocol.

SVFs of iBAT from adult *TCF/Lef:H2B-GFP* male mice were separated and collected as described above. Cells at passage 1 were transduced with *pUltra-hot-LT* LV particles. Two days after transduction, cells were purified by performing FACS to remove mCherry^-^ and Lin^+^ cells. Immortalized cells were subjected to clonal selection through serial limited dilutions. The expression of GFP in cell colonies under pro-adipogenic induction was used as a readout for the accurate establishment of Wnt^+^ (> 95%) and Wnt^-^ (Wnt^+^ = 0%) adipocyte cell lines, respectively.

### *In vitro* pro-adipogenic differentiation of mouse cells

Immortalized cell lines, isolated adipose SVF cells, MEFs, and BMSCs were seeded into plates at the density of 4 x 10^3^ cells/cm^2^ (cell line and SVF) or 1.5 x 10^4^ cells/cm^2^ (BMSCs) in MesenCult^TM^ Expansion medium, and treated with MesenCult^TM^ Adipogenic Differentiation cocktail (STEMCELL, 05507) with 90% cell confluence. Cells were cultured for various days after pro-adipogenic induction prior to being harvested for following experiments.

Adipogenesis was examined by lipid staining. Briefly, differentiated adipocytes were washed twice with PBS, fixed in 4% paraformaldehyde (PFA) for 15 min, and then stained with Oil Red O solution (Sigma-Aldrich, O1391) for 20 min at ambient temperature or with 20 nM BODIPY in PBS for 15 min at 37°C. Subsequently, cells were washed with PBS before imaging.

For inhibition studies, SVF cells or precursor cell lines, 3 days after pro-adipogenic induction, were treated with each of the following molecules for 4 days: DKK1 (100 ng/ml), IWP-2 (5 μM), LF3 (10 and 20 μM, or 50 μM on cell lines), and LY294002 (2 and 14 μM), along with vehicles (PBS or DMSO).

### SiRNA-mediated knockdown experiment

To knockdown *Ctnnb1* in adipocytes, siRNA probes were purchased from IDT (TriFECTa DsiRNA Kit) and transfected into BMSCs that had been under pro-adipogenic induction for two days. Cells at a density of 5 x 10^4^ cells/cm^2^ were plated with 10 nM of a given siRNA dissolved in 1.5% Lipofectamine RNAiMAX (Invitrogen, 13778150) in Opti-MEM I reduced serum medium (Invitrogen, 31985062) and pro-adipogenic differentiation medium (STEMCELL, 05507). The medium was replaced after 24-hr incubation. Time-lapse imaging was carried out to record the real-time change of GFP signal and cell morphology for the next 2 days. At the end, cells were harvested.

### Detection of activated Wnt/β-catenin signaling in adipocytes differentiated from human BMSCs

Primary human bone marrow stromal cells (hBMSCs) from a 29-year-old Caucasian male were purchased from Lifeline Cell Technology (FC-0057 Lot #06333). Lentiviral shuttle vector *pUltra-hot-Tcf/Lef:H2B-GFP* was generated to serve as Wnt/β-catenin signaling reporter, with mCherry as a reporter for successful transduction of hBMSCs. The inserted fragment *Tcf/Lef:H2B-GFP* was amplified from a template plasmid (Addgene, 32610) with primers 5’-agagatccagtttggttaattaatattaaccctcactaaagg -3’ and 5’- tggagccgacacgggttaatttacttgtacagctcgtc -3’. The 2335-bp PCR product was subsequently gel-purified. The vector template *pUltra-hot* (Addgene, 24130) was linearized with PacI and gel-purified. The purified PCR products and linearized vectors were assembled using a GenBuilder Plus kit (Genscript, L00744-10) according to the manufacturer’s instructions. The assembled vector was termed *pUltra-hot-Tcf/Lef:H2B-GFP* as the final shuttle vector. LV production was described as above.

Human BMSCs were plated at a density of 5.0 x 10^4^ cells/cm^2^ in Stemlife MSC-BM Bone Marrow Medium (Lifeline Cell Technology, LL-0026). When cultured cells reached 60-70% confluency, concentrated LV pellets were resuspended and mixed with LentiBlast reagent A and B (1:100) (OZBIOSCIENCES, LBPX500) in a freshly prepared medium and incubated with cells. Forty-eight hrs after viral transduction, hBMSCs were cultured in either pro-adipogenic induction medium (Lifeline Cell Technology, LL-0059) or pro-osteogenic induction cocktail (Lifeline Cell Technology, LM-0023) as positive control for Wnt/β-catenin signaling activity for 7 days, respectively. Cells were subsequently subjected to immunofluorescent assays.

### Immunofluorescence

Adipose tissue was fixed in 4% PFA at 4°C overnight, followed by dehydration through serial ethanol. Samples were processed into paraffin-embedded serial sections at 6 μm. For immunostaining, paraffin-embedded sections were deparaffinized in xylene and subsequently rehydrated. After the incubation of the slides in boiling Tris-EDTA antigen retrieval buffer for 10 min, the tissues were blocked in PBS containing 10% BSA for 60 min, followed by incubation with primary antibodies against GFP (1:500), Perilipin (1:500), UCP1 (1:300), active β-catenin (1:100), Tcf3 (1:500), and Tcf1 (1:100) at 4°C overnight. Slides were then incubated with secondary antibodies (1:500) at room temperature for 60 min. After washing, sections were processed with Autofluorescence Quenching Kit treatment (Vector, SP-8400) for 5 min to remove unspecific fluorescence on sections due to red blood cells and structural elements such as collagen and elastin. Sections were stained with 4’,6-diamidino-2-phenylindole (DAPI) and mounted with mounting medium (Vector, Vibrance^TM^ Antifade). Images of tissue samples were captured using a Nikon confocal Microscope A1 HD25 and analyzed using the ImageJ software (Version 1.51S). For quantification of Wnt^+^ adipocytes, we randomly chose 12-18 images from each tissue of one mouse. Microscopic pictures (10x objective) were taken, and the total cell number and the number of GFP^+^ adipocytes within each image were counted.

For cryosections, samples were dehydrated in 30% sucrose PBS solution overnight at 4°C, embedded in optimal cutting temperature compound (Tissue-Plus; Fisher Healthcare), and frozen by liquid nitrogen for solidification. Embedded samples were cryosectioned (Leica, CM1860) at 8 μm and subjected to immunofluorescent staining with primary antibodies against Tcf4 (1:1:00) and Tcf1 (1:100) as described above.

For immunostaining on cell culture, cells were fixed with 4% PFA for 15 min at ambient temperature and then permeabilized in 0.25% Triton X-100 in PBS for 10 min, followed by blocking with 10% BSA in PBS for 30 min. Cells were then incubated with primary antibodies against GFP (1:500), Perilipin (1:500), Runx2 (1:300), mCherry (1:200), Pparγ (1:400), Adiponectin (1:100), Cyp2e1 (1:200), and Cidea (1:100) at 4°C overnight, respectively. Subsequently, cells were stained with secondary antibodies (1:500) at room temperature for 60 min, followed by counterstaining with DAPI. Images were obtained using a Nikon confocal Microscope A1 HD25 and analyzed using the ImageJ software.

### Western blotting

Proteins were extracted from iWAT or cultured adipocytes using Minute™ Total Protein Extraction Kit (Invent Biotechnologies, AT-022) according to manufacturer’s instructions. 20 μg of proteins were separated by SDS-PAGE (Invitrogen, NW04127BOX) and transferred onto a 0.22 μm Nitrocellulose membrane (LI-COR Biosciences, 926-31092). Membranes were blocked in Tris-buffered saline (TBS) with 0.1% Tween 20 and 5% BSA for 1 hour, followed by overnight incubation with primary antibodies against AKT (1:1,000), p-AKT (1:2,000), GSK-3β (1:1,000), p-GSK-3β (1:500), 4E-BP1 (1:1,000), p-4E-BP1 (1:1,000), UCP1 (1:1,000), OXPHOS cocktail (1:250), or β-actin (1:10,000) in blocking solution at 4°C overnight. Samples were then incubated with secondary antibodies conjugated to IRDye 800 or IRDye 680 (LI-COR Biosciences) diluted at 1:5000 for 1 hour. Immunoreactive protein was detected by Odyssey Imaging System (LI-COR Biosciences).

### RNA preparation and quantitative RT-PCR

Total RNA was extracted from tissue or cells according to protocol for the RNeasy Lipid Tissue Mini Kit (Qiagen, 74804) or RNeasy Mini Kit (Qiagen, 27106) accordingly. Complementary DNA was synthesized using RevertAid First Strand cDNA Synthesis kit (Thermo Scientific, K1622) according to the protocol provided. Quantitative PCR was performed using a C1000 Touch^TM^ thermal cycler (BioRad, CFX96^TM^ Real-Time System). Values of each gene were normalized to reference genes, *36B4* or *Adipoq*, using the comparative Ct method. Primer sequences are provided in Supplementary table 1.

### ScRNA-seq and scATAC-seq

SVF cells isolated from iBAT (3 males and 4 females) and iWAT (10 males and 10 females) of adult *T/L-GFP* mice were cultured and induced to differentiate into adipocytes as described above. Following separation by FACS, approximately 3,000 iBAT-derived GFP^+^, iBAT-derived GFP^-^, iWAT-derived GFP^+^ cells each, and 4,000 iWAT-derived GFP^-^ cells were loaded using Chromium Single Cell 3’ v2 Reagent Kit, and libraries were run on the Illumina NextSeq 550 Sequencing System as 150-bp paired-end reads. Sequencing data were processed on CellRanger v.3.1.0 pipeline with default parameters, including demultiplexing, conversion to FASTQs using bcl2fastq2 v.2.27.1 software, alignment to the mm10 mouse reference genome, filtering, and unique molecular identified (UMI) counting. Gene expression count matrixes were obtained with 2,329 iBAT GFP^+^ cells (32,539 mean reads and 1,313 median genes per cell), 2,666 iBAT GFP^-^ cells (29,534 mean reads and 2,076 median genes per cell), 2,318 iWAT GFP^+^ cells (35,693 mean reads and 2,282 median genes per cell), and 3,137 iWAT GFP^-^ cells (24,360 mean reads and 2,565 median genes per cell). Further scRNA-seq data analysis was performed using the Seurat (Satija et al., 2015) package v.3.2.2. Low-quality cells with fewer than 500 detected genes, doublets with more than 4000 transcripts, and cells with mitochondrial fraction rate higher than 40% were excluded from the analysis. Normalized and scaled data were clustered using the top 20 significant principal components of highly variable genes with the parameter “resolution = 0.7”. Uniform Manifold Approximation and Projection (UMAP) dimensionality reduction was performed to visualize the resulting clusters. To filter out non-adipocytes, preadipocytes, and endothelial cells, cell clusters identified by high expression levels of precursor marker *Pdgfra* (Sun et al., 2020a) and endothelial marker *Cdh5* (Corada et al., 2001) were removed, and *Adipoq*-expressing cells were considered as adipocytes (Lara-Castro et al., 2007) that were reperformed to assign clusters (Figure 3-figure supplement 1A). The final reported datasets consist of 1,710 Wnt^+^ and 827 Wnt^-^ adipocytes for iBAT, 984 Wnt^+^ and 361 Wnt^-^ adipocytes for iWAT, respectively (Figure 3B). Top marker genes of each cluster and library were determined by the *FindAllMarkers* function with the parameters “only.pos = TRUE, min.pct = 0.2, logfc.threshold = 0.25”. Heat maps of top 30 enriched genes from each library was generated using the *DoHeatmap* function (Figure 3D and E). Hallmark gene sets, GO Biological Processes ontology, KEGG pathway, and Wikipathway analyses were performed using the Molecular Signatures Database (MSigDB) (https://www.gsea-msigdb.org/gsea/msigdb) to estimate the functional enrichment and biological pathway of input Wnt^+^ adipocytes based on the differential gene expression (FDR < 0.05) (Figure 3G-J).

For scATAC-seq experiment, SVF cells from adult *T/L-GFP* mice (11 males for iBAT; 7 males and 2 females for iWAT) were induced and adipocytes were acquired as the same workflow for scRNA-seq used (Figure 3A). To isolate nuclei, cells were lysed for 4 min on ice according to 10x genomics protocol (CG000169 Rev D). ScATAC-seq libraries were prepared using the 10x Genomics platform with the Chromium Single Cell ATAC Library & Gel Bead Kit as recommended by the manufacturer and sequenced on the Illumina NextSeq 550 Sequencing System as 150-bp paired-end reads. Peak matrixes and metadata were generated by Cell Ranger ATAC v.1.1.0 pipeline with default parameters and aligned to the mm10 mouse reference genome. Overall, scATAC-seq datasets containing 3,284 iBAT GFP^+^ cell (17,706 median fragments per cell), 4,650 iBAT GFP^-^ cells (9,994 median fragments per cell), 766 iWAT GFP^+^ cells (35,731 median fragments per cell), and 1,034 iWAT GFP^-^ cells (42,083 median fragments per cell) were obtained. Further data analysis was performed using the Signac (Stuart et al., 2019) v.1.0.0 package. In brief, outlier cells with < 2,000 or > 40,000 peaks (iWAT GFP^+^ > 60,000 peaks, iWAT GFP^-^ > 90,000 peaks), < 20% reads in peaks, > 0.025 blacklist ratio, > 4 nucleosome signal or TSS-enrichment < 2, were considered as low-quality cells or doublets and were removed from downstream analyses. Data of each experiment based on the 95% most common feature were then normalized and scaled through the *FindTopFeatures* and *RunSVD* functions. UMAP and k-nearest neighbor (KNN) were applied to perform non-linear dimension reduction using latent semantic indexing (LSI) (Cusanovich et al., 2018) and clustering analysis with the parameter of “resolution = 0.7”. To exclude potential non-adipocytes, gene activity matrix for each experiment was generated by summarizing the accessibility in promoter (TSS and up to 2kb upstream) and cells with high activity of adipocyte specific marker *Adipoq* (Lara-Castro et al., 2007) were retained for further analysis. We merged the datasets of Wnt^+^ and Wnt^-^ adipocytes derived from iBAT and iWAT, respectively, and applied UMAP for the data visualization (Figure 3C, Figure 3-figure supplement 1B). Differential chromatin accessibilities between Wnt^+^ and Wnt^-^ adipocytes were visualized by the *CoveragePlot* function (Figure 3-figure supplement 1C, D), and motif activities were computed through *Chromvar* (Schep et al., 2017) within Signac (Figure 3-figure supplement 1E, F).

### Mitochondrial membrane potential

Mitochondrial membrane potentials of Wnt^+^ adipocytes were measured through MitoTracker Deep Red (far red-fluorescent dye) staining. Briefly, SVF cells derived from iBAT were isolated and collected with the method described above. After 7 days of pro-adipogenic induction, cells were incubated with 100 nM MitoTracker™ Deep Red FM (Invitrogen, M22426) for 30 min at 37°C. After washing twice with PBS, GFP (abs/em ∼ 488/509 nm) fluorescence of differentiated adipocytes and MitoTracker Deep Red (abs/em ∼ 644/665 nm) of active mitochondria were monitored using the Nikon confocal Microscope A1 HD25 and analyzed using the ImageJ software.

### Respiration measurements

Cellular oxygen consumption rate (OCR) was measured using the Seahorse XFe24 analyzer (Agilent Technologies). GFPpos-1, GFPpos-2, GFPneg-1, and mBaSVF cells were seeded in an XF24 cell culture microplate (Agilent Technologies, 102342-100) at a density of 20,000 cells per well. After 5 days of pro-adipogenic induction, 500 μl XF assay medium containing 1 mM pyruvate, 2 mM glutamine, and 10 mM glucose was added to each well. Cells were subjected to a mitochondrial stress test by adding oligomycin (5 μM), FCCP (1.25 μM), and rotenone and antimycin A (5 μM) according to the manufacturer’s instructions (Agilent Technologies, 103015-100).

### Core body temperature measurement

Core body temperatures of *T/L-DTA* mice and control littermates were obtained at RT, 6-, 12-, 24-, 36-, 48-, 60-hour in cold exposure using rector thermometer (Kent Scientific, WD-20250-91) and rectal probe (Kent Scientific, RET-3). Before recording temperatures, measuring instruments were calibrated using ice-water bath each time. Probe was gently inserted into the mouse rectum at least 2 cm. Detailed data see Supplementary table 4.

### Intraperitoneal glucose tolerance test

Mice were fasted for 6 hours starting from 9 am to 3 pm on the testing day Glucose was administered intraperitoneally (2 g/kg body weight) and blood glucose levels were determined from tail vein blood samples using the ACCU-CHEK active glucometer at several time points post glucose injection.

### Cell implantation

Immortalized GFPpos and mBaSVF cells were differentiated under pro-adipogenic conditions, respectively, as described above. After 2-day induction, 1.3 x 10^6^ of each group of committed cells were gently re-suspended and embedded in 110 μl Matrigel (Corning, 356231). The complex was subsequently injected subcutaneously into the left abdomen of C57BL/6J wild-type mice at 8 weeks of age. To trace the fate and the locations of implanted cells, blue agarose beads (150 - 300 μm in diameter; Bio-Rad, 153-7301) were included as an indicator in the implanted complexes. Mice were then kept under regular chow diet at ambient temperature, and ectopically formed fat pads were identified and confirmed by gross and histological examinations 2-week after injection. Accordingly, cell-implanted mice were subject to GTT at 2-week time point after cell implantation.

### Statistics

Statistical analyses were performed using GraphPad Prism 9.0 (GraphPad Software), and Excel (Microsoft). All data were represented as mean ± s.e.m, except where noted. A two-sample unpaired Student’s t-test was used for two-group comparisons. One-way ANOVA followed by Tukey’s test was used for multiple group comparisons, two-way repeated-measures ANOVA followed by Bonferroni’s test was used for Seahorse measurement and GTT results from multiple groups. *P* values below 0.05 were considered significant throughout the study and is presented as \**P* < 0.05, \*\**P* < 0.01, or \*\*\**P* < 0.001.

## Data and code availability

The GEO accession number for the scRNA-seq and scATAC-seq data is GSE164747. Scripts used to process and analyze data in this paper have been deposited to GitHub: https://github.com/ychen-lab/Wnt-positive-adipocyte.

## ACKNOWLEDGEMENTS

We thank members of the Chen Lab for providing technical advice and sharing reagents. We thank Dr. Konrad Basler of the University of Zurich for his kind gift of *Ctnnb1*^dm^ mice. This work was supported by a Carol Lavin Bernick Faculty Grant from Tulane University, the John L. and Mary Wright Ebaugh Endowed Chair Fund, and a grant (R01DK128907) from the NIH to Y.C. T.Y. was supported by an American Heart Association Predoctoral Fellowship (20PRE35040002).

## AUTHOR CONTRIBUTIONS

Z.L., T.C., W.T. and Y.C. conceived the study and designed experiments. D.B. helped to design the initial experiments and provide mentoring and intellectual advice during the study. Z.L., T.C., S.Z., T.Y. performed experiments. Y.G. and H.D. helped to analyze bioinformatic data. Z.L., T.C., S.Z., W.T. and Y.C. collected, analyzed, and interpreted data. Z.L. and T.C. wrote manuscript, and Z.L., T.C., W.T. and Y.C edited the manuscript.

## SUPPLEMENTARY TABLE TITLES (Supplementary table 2-4 are provided as excel files)

**Supplementary table 1.**
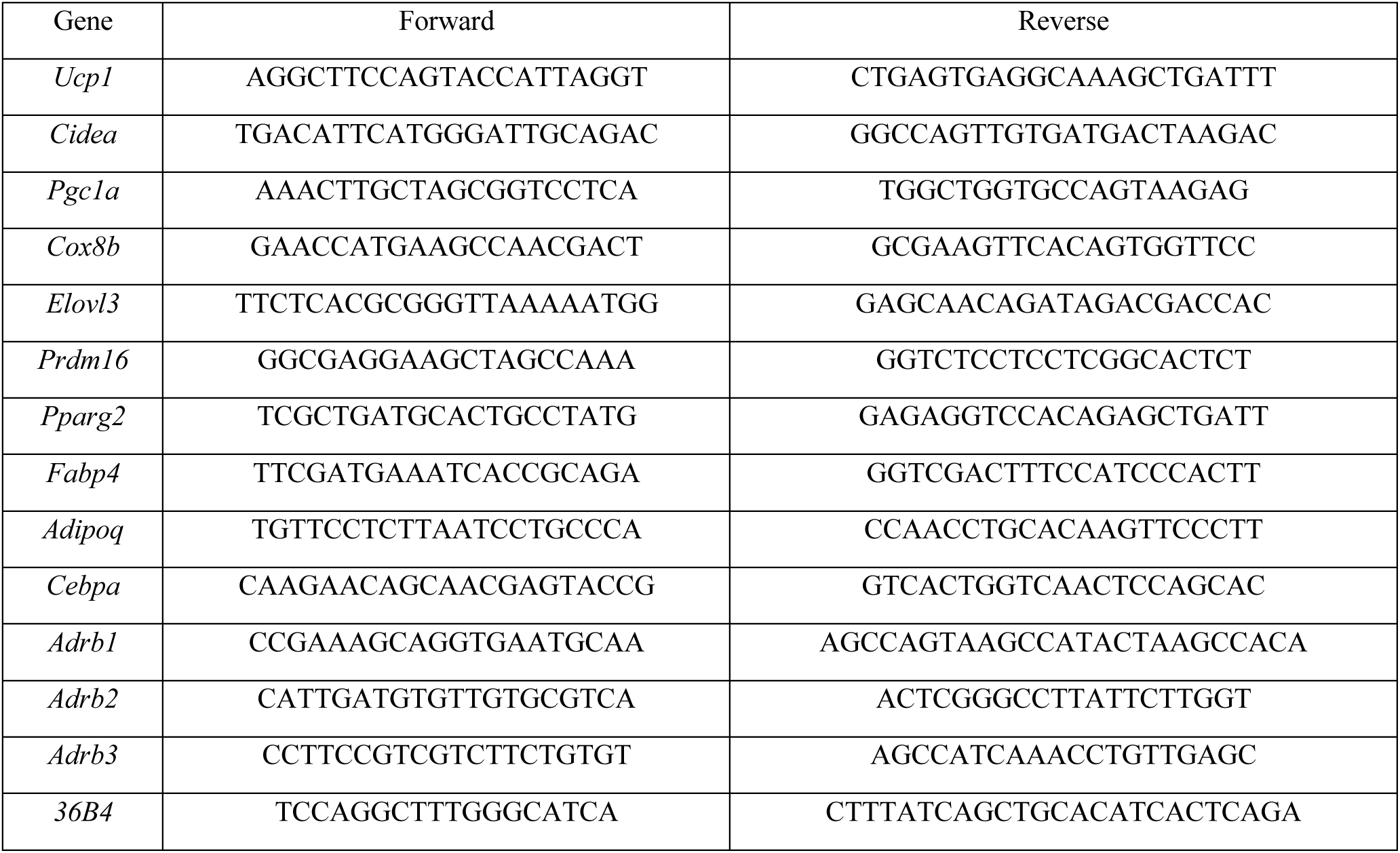
Primer sequences used for qRT-PCR.

**Supplementary table 2.**

Sheet 1: Enriched genes in scRNA-seq.

Sheet 2: Overrepresented DNA motifs in scATAC-seq.

**Supplementary table 3.** Pathway analyses results of DEGs enriched in iWAT-derived Wnt^+^ adipocytes in scRNA-seq.

**Supplementary table 4.** Core body temperature results of *T/L-DTA* and control mice. Core body temperature below 34.5°C was considered as hypothermia and such mice were subsequently euthanized.

---

**Figure 1-figure supplement 1.**
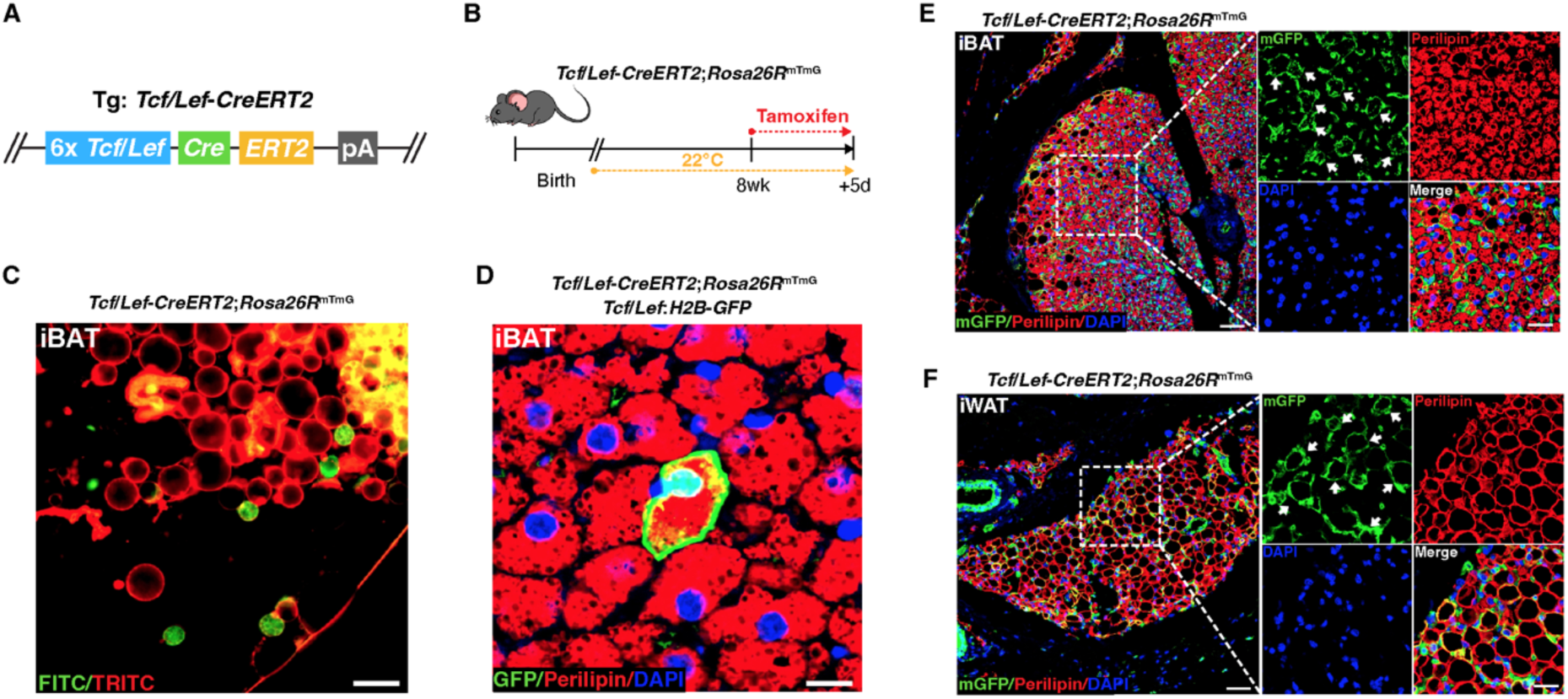
An inducible mouse model validates the existence of Wnt/β-catenin-positive adipocytes. (**A**) Schematic overview of the *TCF/Lef-CreERT2* allele. (**B**) Illustration of the validation experiments in Wnt/β-catenin-positive lineage reporter mice. (**C**) Microscopy images of freshly isolated mGFP-positive adipocytes from iBAT of *TCF/Lef-CreERT2*;*Rosa26R*^mTmG^ mice. (**D**) GFP expression in the cell membrane and nucleus of an adipocyte in *Tcf/Lef-CreERT2*;*Rosa26R*^mTmG^;*T/L-GFP* mice. Scale bar, 10 μm. (**E** and **F**) *Tcf/Lef-CreERT2;Rosa26R*^mTmG^ mice exhibited mGFP-positive adipocytes (white arrows) in iBAT (**E**) and iWAT (**F**) after tamoxifen treatment. *n* = 4 mice each. Scale bars, 50 μm; close-up scale bars, 20 μm. White arrows point to Wnt^+^ adipocytes.

**Figure 1-figure supplement 2.**
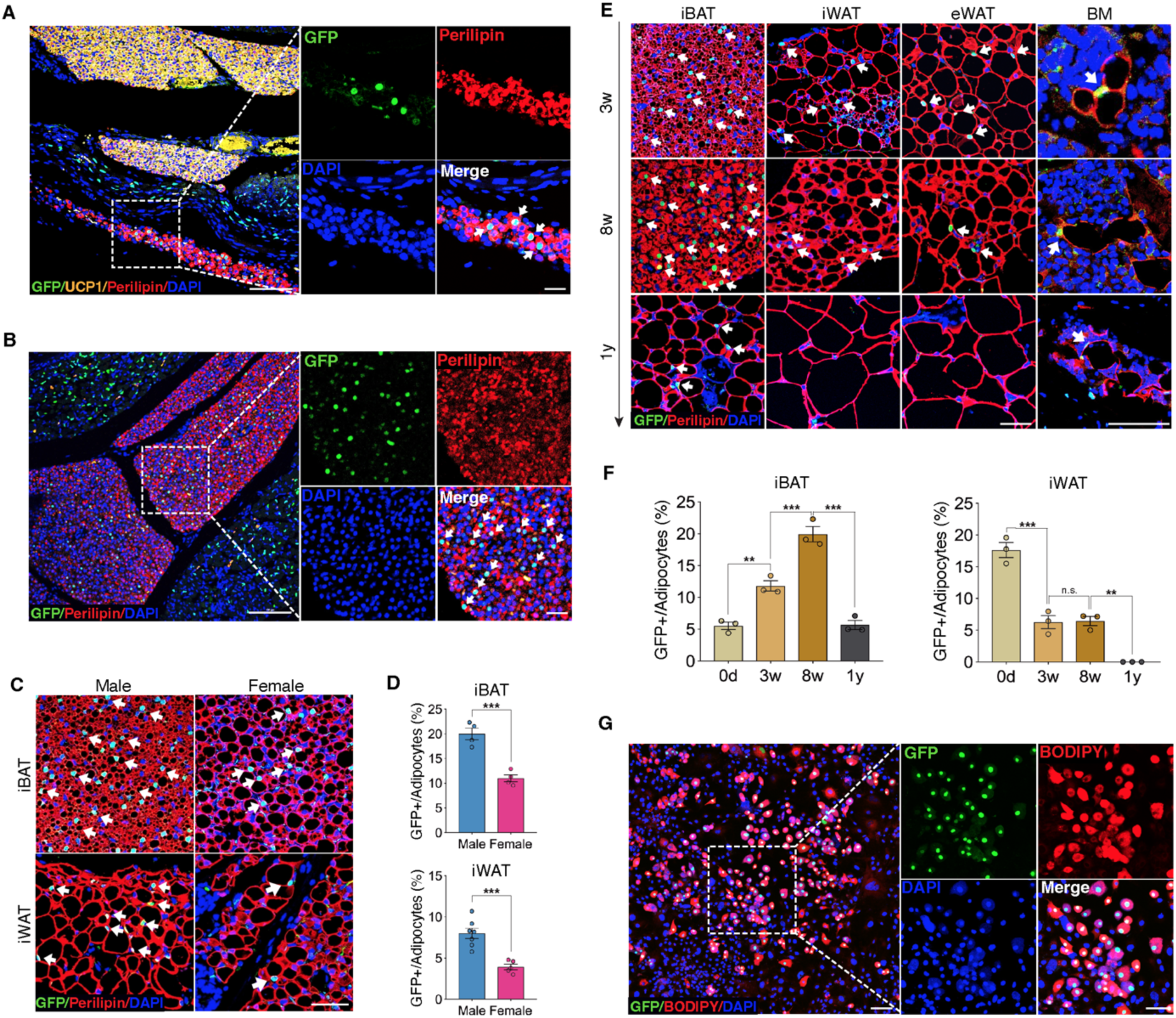
Age- and sex-dependent dynamics of Wnt^+^ adipocytes. (**A**) Immunofluorescent images of subcutaneous WAT adjacent to iBAT from E17.5 *T/L-GFP* mouse showing initial appearance of Wnt^+^ adipocytes. *n* = 3 embryos. Scale bar, 100 μm; close-up scale bar, 25 μm. (**B**) Immunofluorescent images of iBAT from E18.5 *T/L-GFP* mice showing initial appearance of Wnt^+^ adipocytes. *n* = 3 embryos. Scale bar, 100 μm; close-up scale bar, 25 μm. (**C**) Immunofluorescent images of iBAT and iWAT from adult male and female mice showing Wnt^+^ adipocytes. *n* = 4-7 mice. Scale bar, 50 μm. (**D**) Quantification of Wnt^+^ adipocytes among total adipocytes in (**C**). (**E**) Immunofluorescent images of various fat depots from *T/L-GFP* mice at different ages showing the presence of Wnt^+^ adipocytes. *n* = 3 mice each. Scale bar, 50 μm. (**F**) Quantification of Wnt^+^ adipocytes among total adipocytes in (**E**). (**G**) Immunofluorescent images of Wnt^+^ adipocytes induced from E13.5 MEFs *in vitro*. *n* = 2 independent experiments from 8 embryonic mice, 3 independent wells each. Scale bar, 100 μm; close-up scale bar, 20 μm. White arrows in all panels point to Wnt^+^ adipocytes. Data are mean ± s.e.m., \*\**P* < 0.01, \*\*\**P* < 0.001, n.s., not significant, unpaired Student’s t-test (**D**) or one-way ANOVA followed by Tukey’s test (**F**).

**Figure 1-figure supplement 3.**
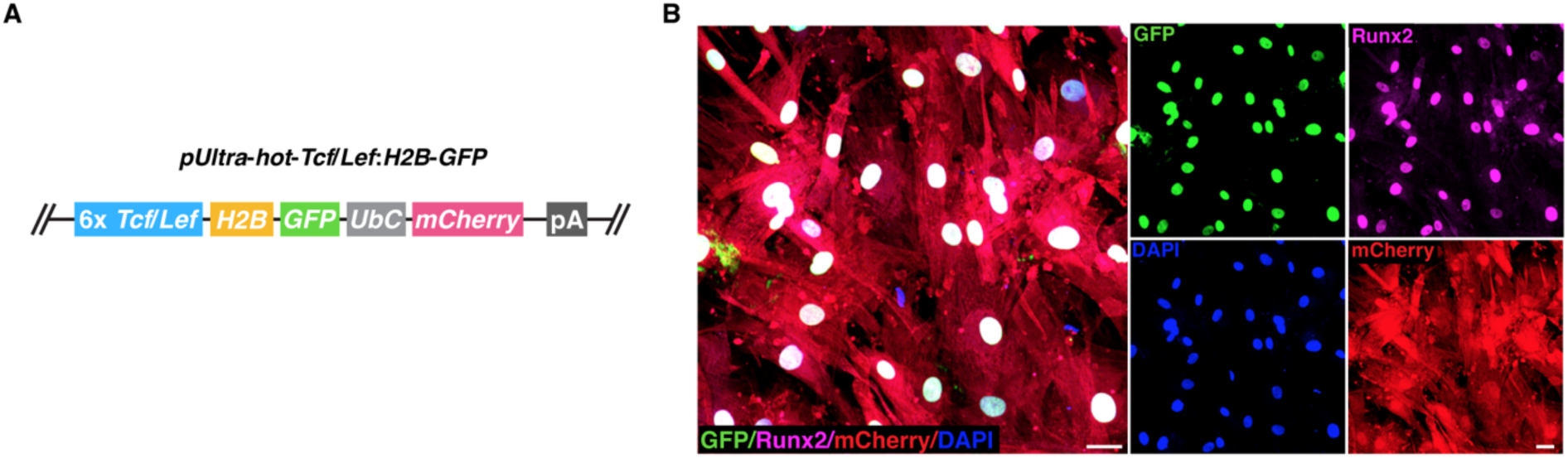
Infection of lentiviral virus in hBMSCs study. (**A**) Construct of the lentiviral shuttle vector. (**B**) Representative immunofluorescent images of infected hBMSCs in pro-osteogenic differentiation medium showing the expression of GFP in the nuclei of osteocytes, indicative of the activation of Wnt/β-catenin signaling. *n* = 2 independent experiments, 3-4 independent wells each. Scale bars, 50 μm.

**Figure 2-figure supplement 1.**
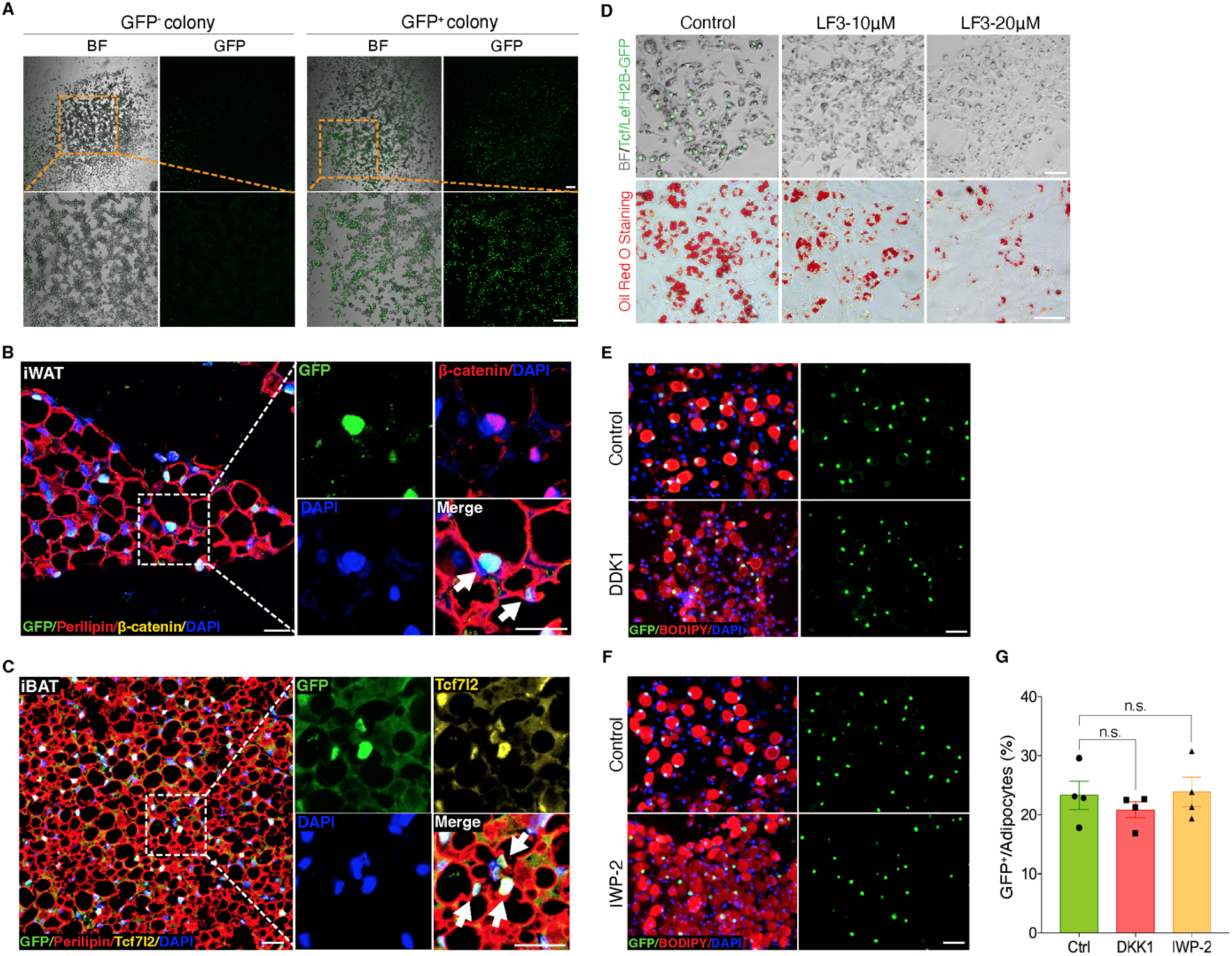
Wnt/β-catenin signaling in adipocytes is activated intracellularly. (**A**) Microscopy images of GFP^-^ and GFP^+^ colonized adipocytes after 7-day differentiation induced from BMSCs of *T/L*-*GFP* mice. *n* = 4 independent experiments. Scale bar, 200 μm. (**B** and **C**) Immunofluorescent images of active β-catenin (**B**) and TCF7L2 (**C**) in Wnt^+^ adipocytes. *n* = 3 mice each. Scale bars, 20 μm (**B**), 25 μm (**C**). (**D**) Microscopy images and Oil Red O staining of Wnt^+^ adipocytes induced from iBAT-derived SVF cells of *T/L*-*GFP* mice with control and LF3 treatment in different doses. *n* = 5 independent experiments. Scale bars, 50 μm. (**E** and **F**) Immunofluorescent images of induced Wnt^+^ adipocytes from BMSCs of *T/L*-GFP mice treated with DKK1 (**D**) and IWP-2 (**E**). *n* = 4 independent experiments each. Scale bars, 50 μm. (**G**) Quantification of Wnt^+^ adipocytes among total adipocytes in (**E**) and (**F**). Data are mean ± s.e.m., n.s., not significant, one-way ANOVA followed by Tukey’s test.

**Figure 2-figure supplement 2.**
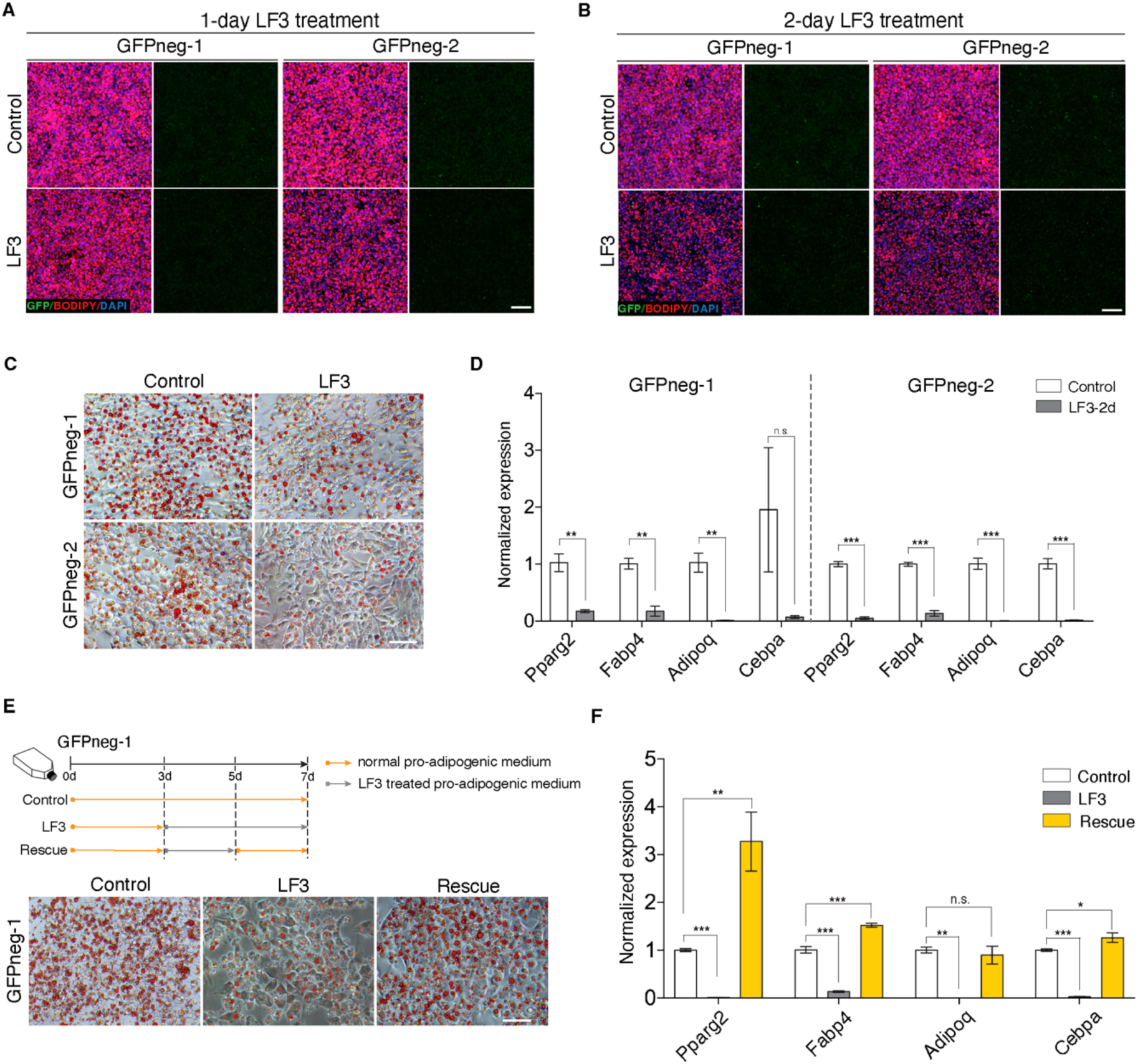
Wnt/β-catenin signaling plays a role in adipogenesis of Wnt^-^ adipocytes. (**A** and **B**) Immunofluorescent images of Wnt^-^ adipocytes induced from two independent cell lines (GFPneg-1 and -2) with LF3 treatment (50 μM) for one (**A**) and two days (**B**), respectively. LF3 was added into the medium after 3-day pro-adipogenic induction. Note that by 1-day LF3 administration, Wnt^-^ adipocytes began to show delayed adipogenic maturation. By 2-day LF3 administration, delayed adipogenic maturation in Wnt^-^ adipocytes became clear, compared to controls. Scale bar, 100 μm. (**C**) Microscopy images of Oil Red O staining in adipocytes induced from GFPneg-1 and -2 cell lines showing reduced lipid accumulation by LF3 treatment for two days. *n* = 3 independent experiments. Scale bar, 50 μm. (**D**) Quantitative RT-PCR analysis of the expression of adipogenic genes in (**C**). The levels of mRNA expression are normalized to that of *36B4*. (**E**) Schematic of rescue experiment and microscopy images of Oil Red O staining in adipocytes induced from GFPneg-1 cell line with LF3 treatment. *n* = 3 independent experiments. Scale bar, 50 μm. (**F**) Quantitative RT-PCR analysis of the expression of adipogenic genes in (**E**). The levels of mRNA expression are normalized to that of *36B4*. Data are mean ± s.e.m., \**P* < 0.05, \*\**P* < 0.01, \*\*\**P* < 0.001, n.s., not significant, one-way ANOVA followed by Tukey’s test (**F**) or unpaired Student’s *t*-test (**D**).

**Figure 3-figure supplement 1.**
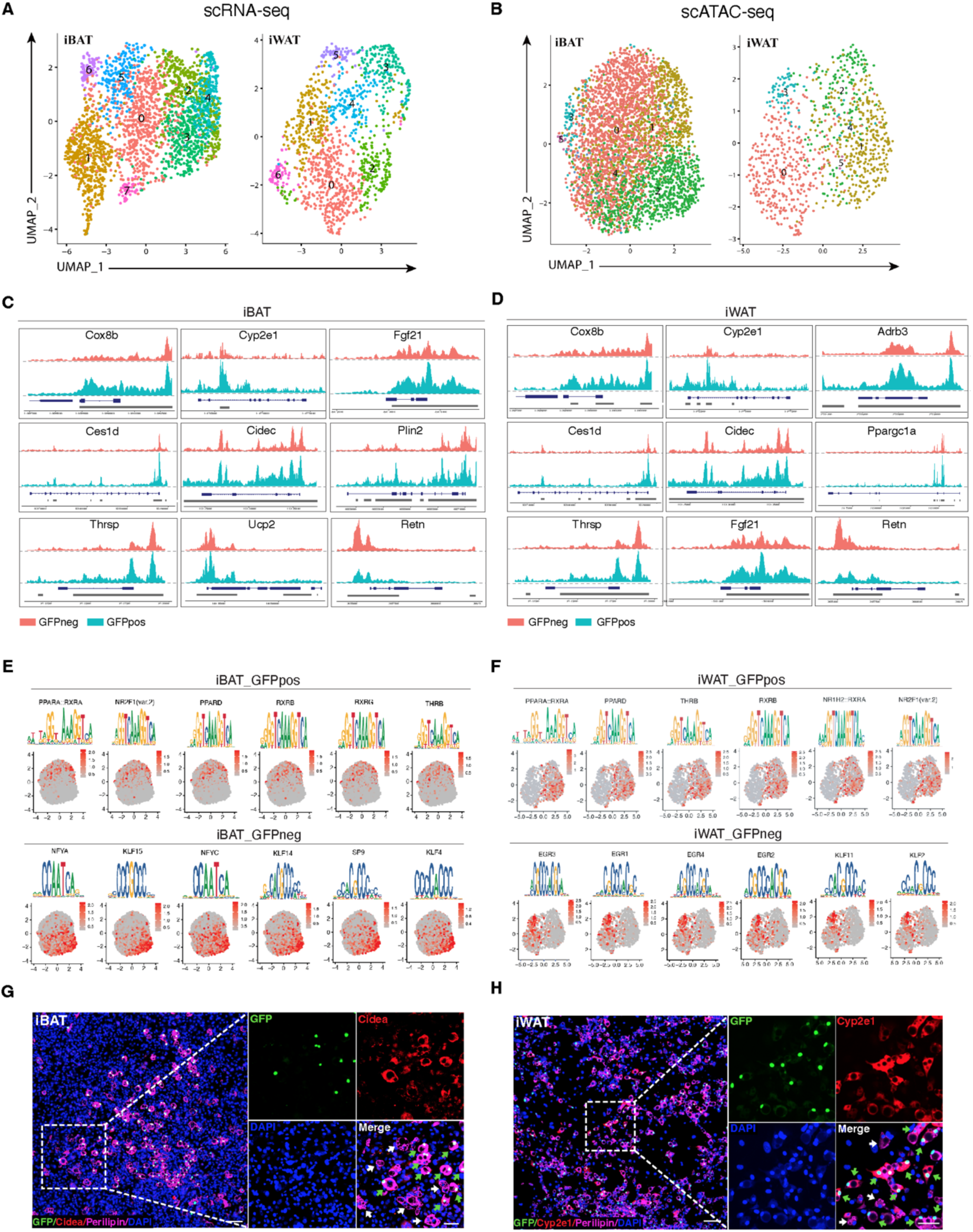
Wnt^+^ adipocytes are distinct from Wnt^-^ ones in molecular signatures. (**A** and **B**) UMAP visualization of unsupervised clustering in scRNA-seq (**A**) and scATAC-seq (**B**). (**C** and **D**) ScATAC-seq tracks of iBAT- (**C**) and iWAT-derived (**D**) Wnt^+^ and Wnt^-^ adipocytes showing distinct chromatin accessibilities of representative genes. (**E** and **F**) Enriched DNA motifs of iBAT- (**E**) and iWAT-derived (**F**) Wnt^+^ (upper panels) and Wnt^-^ (lower panels) adipocytes in scATAC-seq. (**G** and **H**) Immunofluorescent staining of Cidea in adipocytes induced from iBAT-derived SVFs (**G**) and Cyp2e1 in adipocytes induced from iWAT-derived SVFs (**H**). White arrows point to Wnt^-^ adipocytes without Cidea or Cyp2e1 expression, green arrows point to Wnt^+^ adipocytes with positive staining of indicated protein. *n* = 3 independent experiments. Scale bars, 100 μm; close-up scale bars, 25 μm.

**Figure 4-figure supplement 1.**
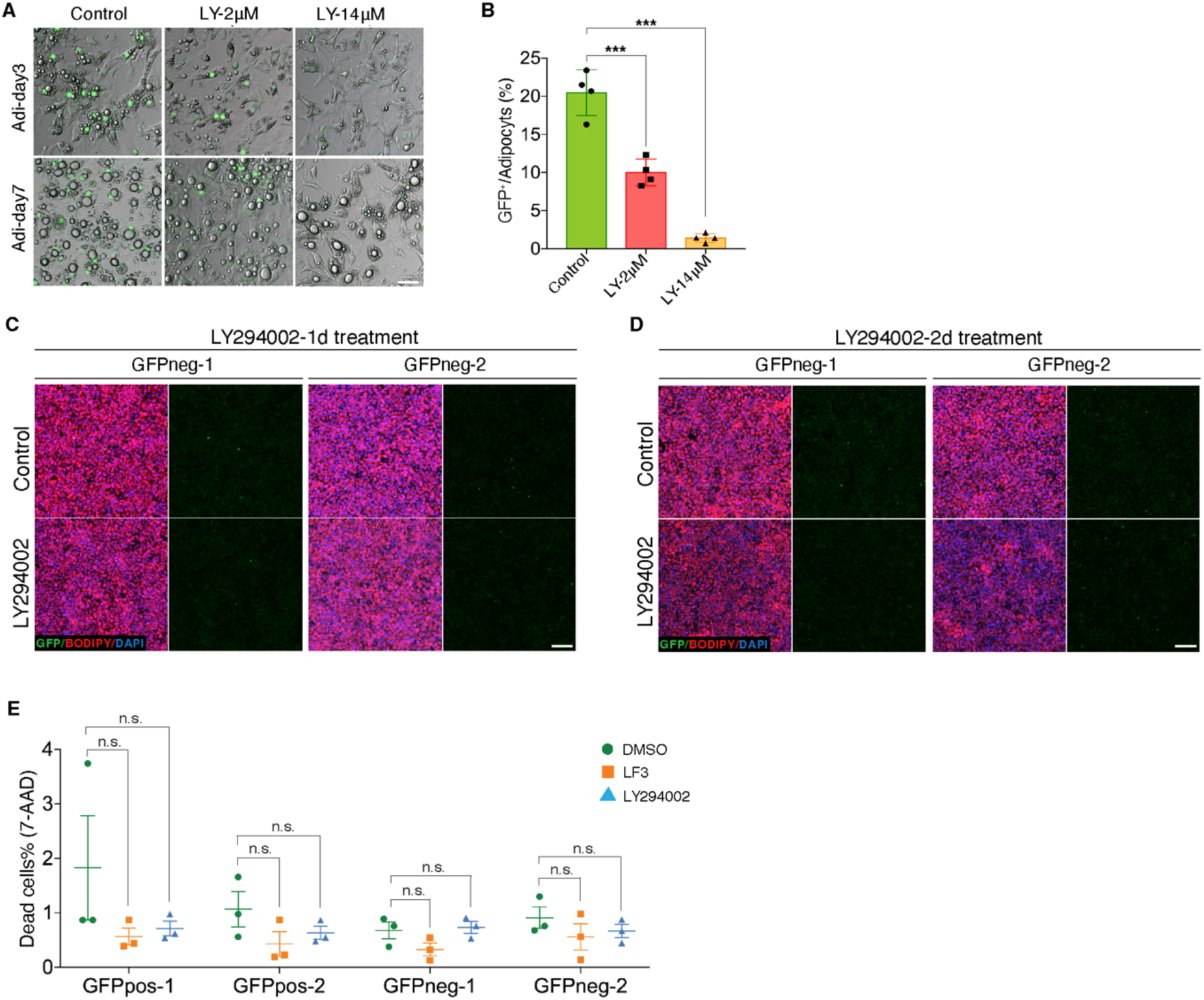
Inhibition of Akt signaling by LY294002 yields similar results as LF3 treatment. (**A**) Microscopy images of LY294002-treated adipocytes showing reduced number of Wnt^+^ adipocytes and delayed adipogenesis after 3- and 7-day treatment compared to controls. *n* = 4 independent experiments. Scale bar, 20 μm. (**B**) Quantification of Wnt^+^ adipocyte among total adipocytes after 7-day treatment in (**A**). (**C** and **D**) Immunofluorescent images of Wnt^-^ adipocytes induced from two independent precursor cell lines (GFPneg-1 and -2) with LY294002 treatment (14 μM) for one (**C**) and two days (**D**), respectively. LY294002 was added into the medium after 3-day pro-adipogenic induction. LY294002 treatment led to slightly reduced lipid storage in Wnt^-^ adipocytes. *n* = 5 independent experiments. Scale bar, 100 μm. (**E**) Quantification of the percentage of 7-AAD-positive cells by flow cytometry analyses showing comparable cell viability in immortalized GFPpos and GFPneg precursor cell lines treated with LF3, LY294002, or DMSO at the same doses. *n* = 3 independent experiments. Data are mean ± s.e.m., \*\*\**P* < 0.001, n.s., not significant, one-way ANOVA followed by Tukey’s test.

**Figure 5-figure supplement 1.**
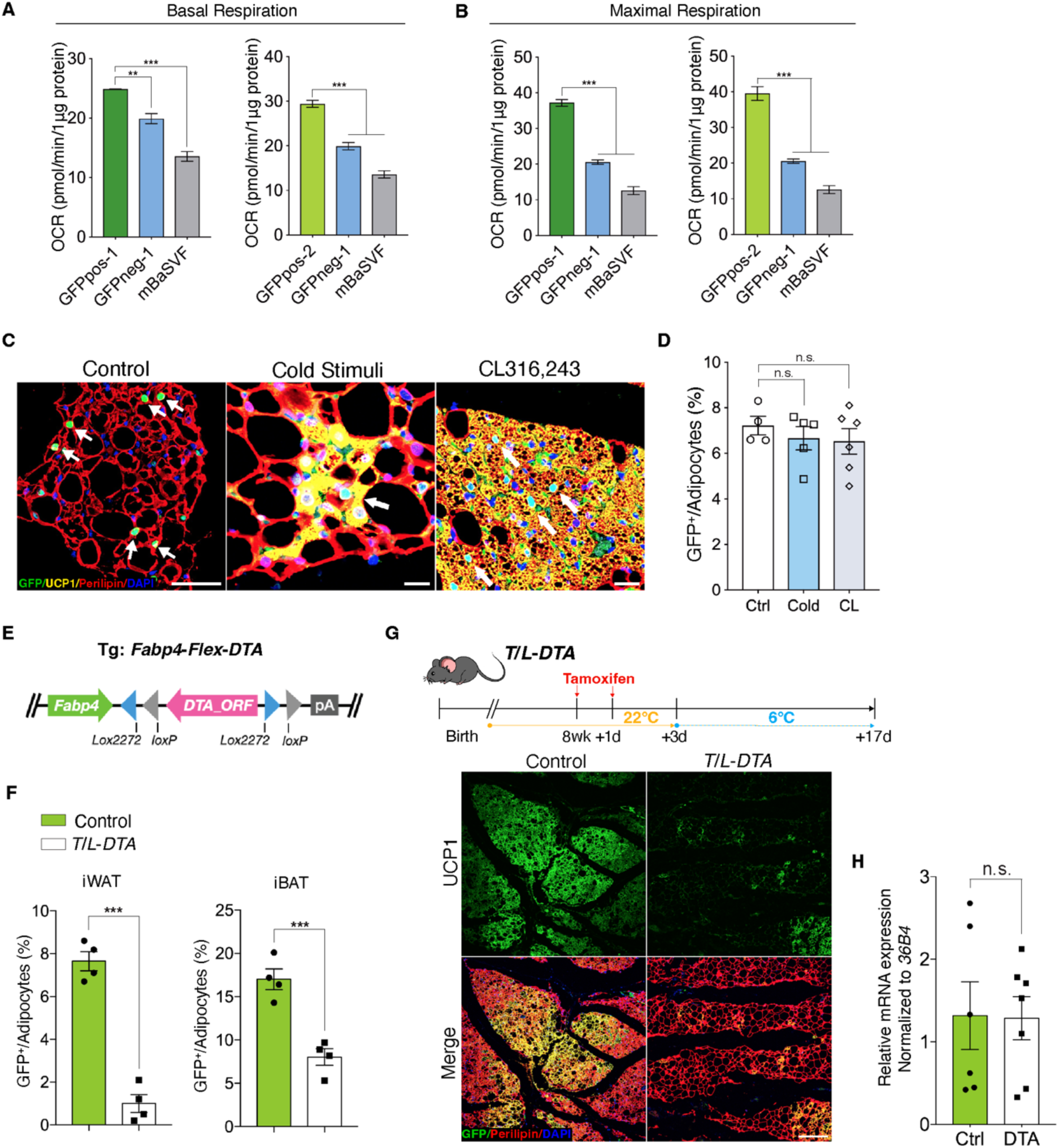
Wnt^+^ adipocytes participate in adaptive thermogenesis in iWAT. (**A** and **B**) Calculated basal (**A**) and maximal (**B**) respiration levels in OCR assay in Figure 5C. (**C**) Immunofluorescent images of iWAT from *T/L-GFP* mice after 2-day cold stress (*n* = 5 mice), CL316,243 treatment (*n* = 6 mice), and control (*n* = 4 mice). Scale bar, 100 μm. White arrows point to Wnt+ adipocytes. (**D**) Quantification of Wnt^+^ adipocytes among total adipocytes in (**C**). (**E**) Schematic overview of the *Fabp4-Flex-DTA* allele. (**F**) Quantification of Wnt^+^ adipocytes among total adipocytes of iWAT and iBAT from *Fabp4-Flex-DTA*;*T/L-GFP* and *T/L*-*DTA*;*T/L-GFP* mice 24 hours after 2-day tamoxifen treatment prior to cold challenge. *n* = 4 mice each. (**G**) Immunofluorescent staining of iWAT from tamoxifen-treated *T/L*-*DTA* mice after 2-week cold exposure. Before cold challenge, mice were rested for 48 hours after the second tamoxifen injection. *n* = 4 mice. Scale bar, 50 μm. (**H**) Quantitative RT-PCR analysis of *Adipoq* expression in iWAT from control (*Fabp4-Flex-DTA*) and *T/L*-*DTA* mice in Fig. 5g, showing comparable results between two groups. *n* = 6 and 7 mice. The levels of mRNA expression are normalized to that of *36B4.* Data are mean ± s.e.m., \*\**P* < 0.01; \*\*\**P* < 0.001, one-way ANOVA followed by Tukey’s test (**A**, **B**, and **D**) or unpaired Student’s *t*-test (**F** and **H**).

## REFERENCES

Bagchi, D.P., Li, Z., Corsa, C.A., Hardij, J., Mori, H., Learman, B.S., Lewis, K.T., Schill, R.L., Romanelli, S.M., and MacDougald, O.A. (2020). Wntless regulates lipogenic gene expression in adipocytes and protects against diet-induced metabolic dysfunction. Mol Metab 39, 100992. 10.1016/j.molmet.2020.100992.

Barbatelli, G., Murano, I., Madsen, L., Hao, Q., Jimenez, M., Kristiansen, K., Giacobino, J.P., De Matteis, R., and Cinti, S. (2010). The emergence of cold-induced brown adipocytes in mouse white fat depots is determined predominantly by white to brown adipocyte transdifferentiation. Am J Physiol Endocrinol Metab 298, E1244–1253. 10.1152/ajpendo.00600.2009.

Bartelt, A., and Heeren, J. (2014). Adipose tissue browning and metabolic health. Nat Rev Endocrinol 10, 24–36. 10.1038/nrendo.2013.204.

Baryawno, N., Sveinbjornsson, B., Eksborg, S., Chen, C.S., Kogner, P., and Johnsen, J.I. (2010). Small-molecule inhibitors of phosphatidylinositol 3-kinase/Akt signaling inhibit Wnt/beta-catenin pathway cross-talk and suppress medulloblastoma growth. Cancer Res 70, 266–276. 10.1158/0008-5472.CAN-09-0578.

Beretta, L., Gingras, A.C., Svitkin, Y.V., Hall, M.N., and Sonenberg, N. (1996). Rapamycin blocks the phosphorylation of 4E-BP1 and inhibits cap-dependent initiation of translation. EMBO J 15, 658–664.

Bostrom, P., Wu, J., Jedrychowski, M.P., Korde, A., Ye, L., Lo, J.C., Rasbach, K.A., Bostrom, E.A., Choi, J.H., Long, J.Z., et al. (2012). A PGC1-alpha-dependent myokine that drives brown-fat-like development of white fat and thermogenesis. Nature 481, 463–468. 10.1038/nature10777.

Cabrae, R., Dubuquoy, C., Cauzac, M., Morzyglod, L., Guilmeau, S., Noblet, B., Feve, B., Postic, C., Burnol, A.F., and Moldes, M. (2020). Insulin activates hepatic Wnt/beta-catenin signaling through stearoyl-CoA desaturase 1 and Porcupine. Sci Rep 10, 5186. 10.1038/s41598-020-61869-4.

Cadigan, K.M., and Liu, Y.I. (2006). Wnt signaling: complexity at the surface. J Cell Sci 119, 395–402. 10.1242/jcs.02826.

Cannon, B., and Nedergaard, J. (2004). Brown adipose tissue: function and physiological significance. Physiol Rev 84, 277–359. 10.1152/physrev.00015.2003.

Chen, M., Lu, P., Ma, Q., Cao, Y., Chen, N., Li, W., Zhao, S., Chen, B., Shi, J., Sun, Y., et al. (2020). CTNNB1/beta-catenin dysfunction contributes to adiposity by regulating the cross-talk of mature adipocytes and preadipocytes. Sci Adv 6, eaax9605. 10.1126/sciadv.aax9605.

Chen, X., Ayala, I., Shannon, C., Fourcaudot, M., Acharya, N.K., Jenkinson, C.P., Heikkinen, S., and Norton, L. (2018). The Diabetes Gene and Wnt Pathway Effector TCF7L2 Regulates Adipocyte Development and Function. Diabetes 67, 554–568. 10.2337/db17-0318.

Cinti, S. (2005). The adipose organ. Prostaglandins Leukot Essent Fatty Acids 73, 9–15. 10.1016/j.plefa.2005.04.010.

Corada, M., Liao, F., Lindgren, M., Lampugnani, M.G., Breviario, F., Frank, R., Muller, W.A., Hicklin, D.J., Bohlen, P., and Dejana, E. (2001). Monoclonal antibodies directed to different regions of vascular endothelial cadherin extracellular domain affect adhesion and clustering of the protein and modulate endothelial permeability. Blood 97, 1679–1684. 10.1182/blood.v97.6.1679.

Crane, J.D., Palanivel, R., Mottillo, E.P., Bujak, A.L., Wang, H., Ford, R.J., Collins, A., Blumer, R.M., Fullerton, M.D., Yabut, J.M., et al. (2015). Inhibiting peripheral serotonin synthesis reduces obesity and metabolic dysfunction by promoting brown adipose tissue thermogenesis. Nat Med 21, 166–172. 10.1038/nm.3766.

Cusanovich, D.A., Hill, A.J., Aghamirzaie, D., Daza, R.M., Pliner, H.A., Berletch, J.B., Filippova, G.N., Huang, X., Christiansen, L., DeWitt, W.S., et al. (2018). A Single-Cell Atlas of In Vivo Mammalian Chromatin Accessibility. Cell 174, 1309–1324 e1318. 10.1016/j.cell.2018.06.052.

Dale, T.C. (1998). Signal transduction by the Wnt family of ligands. Biochem J 329 ( *Pt 2*), 209–223. 10.1042/bj3290209.

Fang, D., Hawke, D., Zheng, Y., Xia, Y., Meisenhelder, J., Nika, H., Mills, G.B., Kobayashi, R., Hunter, T., and Lu, Z. (2007). Phosphorylation of beta-catenin by AKT promotes beta-catenin transcriptional activity. J Biol Chem 282, 11221–11229. 10.1074/jbc.M611871200.

Fang, L., Zhu, Q., Neuenschwander, M., Specker, E., Wulf-Goldenberg, A., Weis, W.I., von Kries, J.P., and Birchmeier, W. (2016). A Small-Molecule Antagonist of the beta-Catenin/TCF4 Interaction Blocks the Self-Renewal of Cancer Stem Cells and Suppresses Tumorigenesis. Cancer Res 76, 891–901. 10.1158/0008-5472.CAN-15-1519.

Ferrer-Vaquer, A., Piliszek, A., Tian, G., Aho, R.J., Dufort, D., and Hadjantonakis, A.K. (2010). A sensitive and bright single-cell resolution live imaging reporter of Wnt/ss-catenin signaling in the mouse. BMC Dev Biol 10, 121. 10.1186/1471-213X-10-121.

Geoghegan, G., Simcox, J., Seldin, M.M., Parnell, T.J., Stubben, C., Just, S., Begaye, L., Lusis, A.J., and Villanueva, C.J. (2019). Targeted deletion of Tcf7l2 in adipocytes promotes adipocyte hypertrophy and impaired glucose metabolism. Mol Metab 24, 44–63. 10.1016/j.molmet.2019.03.003.

Harms, M., and Seale, P. (2013). Brown and beige fat: development, function and therapeutic potential. Nat Med 19, 1252–1263. 10.1038/nm.3361.

Hasegawa, Y., Ikeda, K., Chen, Y., Alba, D.L., Stifler, D., Shinoda, K., Hosono, T., Maretich, P., Yang, Y., Ishigaki, Y., et al. (2018). Repression of Adipose Tissue Fibrosis through a PRDM16-GTF2IRD1 Complex Improves Systemic Glucose Homeostasis. Cell Metab 27, 180–194 e186. 10.1016/j.cmet.2017.12.005.

Hepler, C., Shan, B., Zhang, Q., Henry, G.H., Shao, M., Vishvanath, L., Ghaben, A.L., Mobley, A.B., Strand, D., Hon, G.C., and Gupta, R.K. (2018). Identification of functionally distinct fibro-inflammatory and adipogenic stromal subpopulations in visceral adipose tissue of adult mice. Elife 7. 10.7554/eLife.39636.

Hepler, C., Vishvanath, L., and Gupta, R.K. (2017). Sorting out adipocyte precursors and their role in physiology and disease. Genes Dev 31, 127–140. 10.1101/gad.293704.116.

Hermida, M.A., Dinesh Kumar, J., and Leslie, N.R. (2017). GSK3 and its interactions with the PI3K/AKT/mTOR signalling network. Adv Biol Regul 65, 5–15. 10.1016/j.jbior.2017.06.003.

Ikeda, K., Kang, Q., Yoneshiro, T., Camporez, J.P., Maki, H., Homma, M., Shinoda, K., Chen, Y., Lu, X., Maretich, P., et al. (2017). UCP1-independent signaling involving SERCA2b-mediated calcium cycling regulates beige fat thermogenesis and systemic glucose homeostasis. Nat Med 23, 1454–1465. 10.1038/nm.4429.

Ikeda, K., Maretich, P., and Kajimura, S. (2018). The Common and Distinct Features of Brown and Beige Adipocytes. Trends Endocrinol Metab 29, 191–200. 10.1016/j.tem.2018.01.001.

Ishibashi, J., and Seale, P. (2010). Medicine. Beige can be slimming. Science 328, 1113–1114. 10.1126/science.1190816.

Lara-Castro, C., Fu, Y., Chung, B.H., and Garvey, W.T. (2007). Adiponectin and the metabolic syndrome: mechanisms mediating risk for metabolic and cardiovascular disease. Curr Opin Lipidol 18, 263–270. 10.1097/MOL.0b013e32814a645f.

Longo, K.A., Wright, W.S., Kang, S., Gerin, I., Chiang, S.H., Lucas, P.C., Opp, M.R., and MacDougald, O.A. (2004). Wnt10b inhibits development of white and brown adipose tissues. J Biol Chem 279, 35503–35509. 10.1074/jbc.M402937200.

Lowell, B.B., V, S.S., Hamann, A., Lawitts, J.A., Himms-Hagen, J., Boyer, B.B., Kozak, L.P., and Flier, J.S. (1993). Development of obesity in transgenic mice after genetic ablation of brown adipose tissue. Nature 366, 740–742. 10.1038/366740a0.

Nedergaard, J., and Cannon, B. (2014). The browning of white adipose tissue: some burning issues. Cell Metab 20, 396–407. 10.1016/j.cmet.2014.07.005.

Ng, S.S., Mahmoudi, T., Danenberg, E., Bejaoui, I., de Lau, W., Korswagen, H.C., Schutte, M., and Clevers, H. (2009). Phosphatidylinositol 3-kinase signaling does not activate the wnt cascade. J Biol Chem 284, 35308–35313. 10.1074/jbc.M109.078261.

Oguri, Y., Shinoda, K., Kim, H., Alba, D.L., Bolus, W.R., Wang, Q., Brown, Z., Pradhan, R.N., Tajima, K., Yoneshiro, T., et al. (2020). CD81 Controls Beige Fat Progenitor Cell Growth and Energy Balance via FAK Signaling. Cell 182, 563–577 e520. 10.1016/j.cell.2020.06.021.

Palsgaard, J., Emanuelli, B., Winnay, J.N., Sumara, G., Karsenty, G., and Kahn, C.R. (2012). Cross-talk between insulin and Wnt signaling in preadipocytes: role of Wnt co-receptor low density lipoprotein receptor-related protein-5 (LRP5). J Biol Chem 287, 12016–12026. 10.1074/jbc.M111.337048.

Paulo, E., and Wang, B. (2019). Towards a Better Understanding of Beige Adipocyte Plasticity. Cells 8. 10.3390/cells8121552.

Petrovic, N., Walden, T.B., Shabalina, I.G., Timmons, J.A., Cannon, B., and Nedergaard, J. (2010). Chronic peroxisome proliferator-activated receptor gamma (PPARgamma) activation of epididymally derived white adipocyte cultures reveals a population of thermogenically competent, UCP1-containing adipocytes molecularly distinct from classic brown adipocytes. J Biol Chem 285, 7153–7164. 10.1074/jbc.M109.053942.

Prossomariti, A., Piazzi, G., Alquati, C., and Ricciardiello, L. (2020). Are Wnt/beta-Catenin and PI3K/AKT/mTORC1 Distinct Pathways in Colorectal Cancer? Cell Mol Gastroenterol Hepatol 10, 491–506. 10.1016/j.jcmgh.2020.04.007.

Rajbhandari, P., Arneson, D., Hart, S.K., Ahn, I.S., Diamante, G., Santos, L.C., Zaghari, N., Feng, A.C., Thomas, B.J., Vergnes, L., et al. (2019). Single cell analysis reveals immune cell-adipocyte crosstalk regulating the transcription of thermogenic adipocytes. Elife 8. 10.7554/eLife.49501.

Regard, J.B., Cherman, N., Palmer, D., Kuznetsov, S.A., Celi, F.S., Guettier, J.M., Chen, M., Bhattacharyya, N., Wess, J., Coughlin, S.R., et al. (2011). Wnt/beta-catenin signaling is differentially regulated by Galpha proteins and contributes to fibrous dysplasia. Proc Natl Acad Sci U S A 108, 20101–20106. 10.1073/pnas.1114656108.

Rosen, E.D., and Spiegelman, B.M. (2014). What we talk about when we talk about fat. Cell 156, 20–44. 10.1016/j.cell.2013.12.012.

Rosenwald, M., and Wolfrum, C. (2014). The origin and definition of brite versus white and classical brown adipocytes. Adipocyte 3, 4–9. 10.4161/adip.26232.

Ross, S.E., Hemati, N., Longo, K.A., Bennett, C.N., Lucas, P.C., Erickson, R.L., and MacDougald, O.A. (2000). Inhibition of adipogenesis by Wnt signaling. Science 289, 950–953. 10.1126/science.289.5481.950.

Sarvari, A.K., Van Hauwaert, E.L., Markussen, L.K., Gammelmark, E., Marcher, A.B., Ebbesen, M.F., Nielsen, R., Brewer, J.R., Madsen, J.G.S., and Mandrup, S. (2020). Plasticity of Epididymal Adipose Tissue in Response to Diet-Induced Obesity at Single-Nucleus Resolution. Cell Metab. 10.1016/j.cmet.2020.12.004.

Satija, R., Farrell, J.A., Gennert, D., Schier, A.F., and Regev, A. (2015). Spatial reconstruction of single-cell gene expression data. Nat Biotechnol 33, 495–502. 10.1038/nbt.3192.

Schakman, O., Kalista, S., Bertrand, L., Lause, P., Verniers, J., Ketelslegers, J.M., and Thissen, J.P. (2008). Role of Akt/GSK-3beta/beta-catenin transduction pathway in the muscle anti-atrophy action of insulin-like growth factor-I in glucocorticoid-treated rats. Endocrinology 149, 3900–3908. 10.1210/en.2008-0439.

Schep, A.N., Wu, B., Buenrostro, J.D., and Greenleaf, W.J. (2017). chromVAR: inferring transcription-factor-associated accessibility from single-cell epigenomic data. Nat Methods 14, 975–978. 10.1038/nmeth.4401.

Schwalie, P.C., Dong, H., Zachara, M., Russeil, J., Alpern, D., Akchiche, N., Caprara, C., Sun, W., Schlaudraff, K.U., Soldati, G., et al. (2018). A stromal cell population that inhibits adipogenesis in mammalian fat depots. Nature 559, 103–108. 10.1038/s41586-018-0226-8.

Shao, M., Wang, Q.A., Song, A., Vishvanath, L., Busbuso, N.C., Scherer, P.E., and Gupta, R.K. (2019). Cellular Origins of Beige Fat Cells Revisited. Diabetes 68, 1874–1885. 10.2337/db19-0308.

Stuart, T., Butler, A., Hoffman, P., Hafemeister, C., Papalexi, E., Mauck, W.M., 3rd, Hao, Y., Stoeckius, M., Smibert, P., and Satija, R. (2019). Comprehensive Integration of Single-Cell Data. Cell 177, 1888–1902 e1821. 10.1016/j.cell.2019.05.031.

Sun, C., Sakashita, H., Kim, J., Tang, Z., Upchurch, G.M., Yao, L., Berry, W.L., Griffin, T.M., and Olson, L.E. (2020a). Mosaic Mutant Analysis Identifies PDGFRalpha/PDGFRbeta as Negative Regulators of Adipogenesis. Cell Stem Cell 26, 707–721 e705. 10.1016/j.stem.2020.03.004.

Sun, W., Dong, H., Balaz, M., Slyper, M., Drokhlyansky, E., Colleluori, G., Giordano, A., Kovanicova, Z., Stefanicka, P., Balazova, L., et al. (2020b). snRNA-seq reveals a subpopulation of adipocytes that regulates thermogenesis. Nature 587, 98–102. 10.1038/s41586-020-2856-x.

Tseng, Y.H., Cypess, A.M., and Kahn, C.R. (2010). Cellular bioenergetics as a target for obesity therapy. Nat Rev Drug Discov 9, 465–482. 10.1038/nrd3138.

Valenta, T., Gay, M., Steiner, S., Draganova, K., Zemke, M., Hoffmans, R., Cinelli, P., Aguet, M., Sommer, L., and Basler, K. (2011). Probing transcription-specific outputs of beta-catenin in vivo. Genes Dev 25, 2631–2643. 10.1101/gad.181289.111.

Voight, B.F., Scott, L.J., Steinthorsdottir, V., Morris, A.P., Dina, C., Welch, R.P., Zeggini, E., Huth, C., Aulchenko, Y.S., Thorleifsson, G., et al. (2010). Twelve type 2 diabetes susceptibility loci identified through large-scale association analysis. Nat Genet 42, 579–589. 10.1038/ng.609.

Waki, H., Park, K.W., Mitro, N., Pei, L., Damoiseaux, R., Wilpitz, D.C., Reue, K., Saez, E., and Tontonoz, P. (2007). The small molecule harmine is an antidiabetic cell-type-specific regulator of PPARgamma expression. Cell Metab 5, 357–370. 10.1016/j.cmet.2007.03.010.

Wang, Q.A., Tao, C., Gupta, R.K., and Scherer, P.E. (2013). Tracking adipogenesis during white adipose tissue development, expansion and regeneration. Nat Med 19, 1338–1344. 10.1038/nm.3324.

Wu, J., Bostrom, P., Sparks, L.M., Ye, L., Choi, J.H., Giang, A.H., Khandekar, M., Virtanen, K.A., Nuutila, P., Schaart, G., et al. (2012). Beige adipocytes are a distinct type of thermogenic fat cell in mouse and human. Cell 150, 366–376. 10.1016/j.cell.2012.05.016.

Wu, Y., Kinnebrew, M.A., Kutyavin, V.I., and Chawla, A. (2020). Distinct signaling and transcriptional pathways regulate peri-weaning development and cold-induced recruitment of beige adipocytes. Proc Natl Acad Sci U S A 117, 6883–6889. 10.1073/pnas.1920419117.

Yoshioka, M., Kayo, T., Ikeda, T., and Koizumi, A. (1997). A novel locus, Mody4, distal to D7Mit189 on chromosome 7 determines early-onset NIDDM in nonobese C57BL/6 (Akita) mutant mice. Diabetes 46, 887–894. 10.2337/diab.46.5.887.

Zhang, J., Shemezis, J.R., McQuinn, E.R., Wang, J., Sverdlov, M., and Chenn, A. (2013). AKT activation by N-cadherin regulates beta-catenin signaling and neuronal differentiation during cortical development. Neural Dev 8, 7. 10.1186/1749-8104-8-7.

